# SUMOylation differentially regulates SMCHD1 complex formation and function in a genomic context-specific manner

**DOI:** 10.1101/2024.10.13.618066

**Authors:** Remko Goossens, Mara S. Tihaya, Iris M. Willemsen, Kirsten R. Straasheijm, Patrick J. van der Vliet, Roman González Prieto, Alfred C.O. Vertegaal, Stephen J. Tapscott, Judit Balog, Silvère M. van der Maarel

**Author notes:** Correspondence to, S.M. van der Maarel, Ph.D. Leiden University Medical Center Department of Human Genetics Postal zone S-04-P, 2333 ZA Leiden, The Netherlands. Authors Goossens and Tihaya contributed equally to this manuscript.

## Abstract

Structural Maintenance of Chromosomes Hinge Domain Containing 1 (SMCHD1) is a chromatin repressor regulating gene expression and chromatin architecture of distinct autosomal and X-chromosomal loci. SMCHD1 mutations cause derepression of the D4Z4 macrosatellite repeat-embedded *DUX4* gene in skeletal muscle of facioscapulohumeral muscular dystrophy (FSHD) patients Little is known about the regulation and post-translational modification of SMCHD1. Here we report the SUMOylation dynamics of SMCHD1 and its impact on autosomal single copy and repetitive loci, and the inactive X chromosome (Xi).

We identify that SMCHD1 is SUMOylated primarily at lysine 1374, uncover factors regulating SMCHD1 SUMOylation, and demonstrate that SMCHD1 interacts with chromatin repressors TRIM28, HNRNPK and SETDB1 in a SUMO-dependent manner. We find that SUMOylation impacts Xi engagement of SMCHD1, maintenance of a repressive D4Z4 chromatin structure preventing DUX4 expression, and regulation of *LRIF1* promoter activity. The rapid, SUMO-dependent upregulation of *DUX4* could explain the bursts of DUX4 expression typical for FSHD muscle.

## Introduction

Post-translational modification (PTM) of proteins is a highly dynamic and complex regulatory mechanism affecting many protein features, including protein function and turnover. PTMs include, amongst others, protein phosphorylation, ubiquitination and SUMOylation. As such, PTMs lead to a highly diverse and responsive proteome.

Small Ubiquitin-like modifier (SUMO) is a PTM affecting the epsilon (ε)-amino group of the side chains of specific lysine (K) residues of SUMO protein substrates. Like ubiquitination, SUMOylation consists of the covalent linking of a small polypeptide, which can modify the activity or fate of the substrate. The effects of SUMOylation are substrate-specific and may include subcellular localization, transcriptional activity, protein interactions and protein stability ^1,2^. Three main SUMO proteins have been described in mammals: SUMO1, SUMO2 and SUMO3. SUMO1 was first identified, showing ∼50% homology to SUMO2 and SUMO3, which are identical except for a few amino acids and are often referred to as SUMO2/3 ^3,4^. While SUMOylated lysines are usually located within a ΨKxE SUMO consensus motif, it is challenging to predict SUMO sites in protein sequences ^3,5^. The fraction of SUMOylated substrate at any time can be low, while loss of SUMOylation sites does not always lead to apparent phenotypes, challenging SUMO detection and functional analysis ^1,2^. SUMO-conjugated substrates are predominantly nuclear, where they play a role in regulating various functions ^1,2^. SUMO proteins are partially complementary, although the knock-out of SUMO2 in mice is lethal^4^.

In humans, mutations in Structural Maintenance of Chromosomes Hinge Domain 1 (SMCHD1) can cause Facioscapulohumeral dystrophy (FSHD), a progressive muscle disease leading to weakness and wasting of the facial and upper extremity muscles or Bosma arhinia microphthalmia syndrome (BAMS) ^6–8^, a rare developmental condition characterized by abnormalities of the eyes and nose, as well as hypogonadotropic hypogonadism. It is currently not entirely clear how mutations in SMCHD1 can cause such disparate phenotypes, although some studies suggest a different functional outcome of FSHD and BAMS mutations in SMCHD1 ^9–11^. Nevertheless, the co-existence of identically affected SMCHD1 amino acids in both disorders argues against a simple different functional mutation outcome model for both disorders ^10,12^. SMCHD1 is a chromatin repressor that acts on distinct autosomal loci amongst which homeotic (*Hox*) and protocadherin (*Pcdh*) gene clusters and imprinted loci ^13,14^, as well as being involved in X-chromosome inactivation in females ^15–18^. It also co-ordinates the promoter activity of *LRIF1* encoding an interaction partner of SMCHD1 ^19^. In the context of FSHD, SMCHD1 is essential for the somatic repression of the cleavage stage and germline transcription factor DUX4 expressed from the D4Z4 macrosatellite repeat in the subtelomere of chromosome 4q^20^. The mouse homolog of *DUX4*, *Dux*, which is also embedded in a macrosatellite repeat, has been shown to be regulated in a SUMO-dependent fashion in mouse embryonic stem (ES) cells ^21^.

Two independent studies identified SMCHD1 as a prominent SUMO substrate: one study through anti-hemagglutinin (HA) affinity purification on brain tissue of His6-HA-SUMO1 knock in mice ^3^ and the second study through a SUMO2 global SUMOylation screen in HeLa cells ^5^. While several studies have addressed the function of SMCHD1, often in relation to X inactivation ^22,23^, gene cluster regulation ^13,24^ and higher order chromatin structure ^16–18^, the regulation of SMCHD1 protein activity itself is less intensively studied. This is particularly important as evidence from myogenic differentiation studies suggests a dynamic regulation of SMCHD1 ^25^, which may in part explain the sensitivity of muscle to SMCHD1 dysfunction.

Here, we investigated the role of SUMOylation in the regulation of DUX4 expression in the context of FSHD pathogenesis and other loci regulated by SMCHD1 such as the inactive X chromosome (Xi) and the *LRIF1* promoter. We first studied the global regulation of D4Z4 by SUMOylation in somatic cells, and next addressed the contribution of SMCHD1 SUMOylation to D4Z4 repression and other SMCHD1-regulated loci. We observed that D4Z4 repression in somatic cells is regulated by SUMOylation and report that SMCHD1 can be post-translationally modified by SUMOylation. Through a SUMO-dependent SMCHD1-interactome analysis by mass spectrometry, we identified previously undescribed interaction partners of SMCHD1, such as TRIM28 (KAP1) and HNRNPK, with TRIM28 functioning as an E3 ligase for SMCHD1, while systematic knockdown of all SUMO proteases identified SENP5 as the major SMCHD1 SUMO protease. The SUMO-dependent SMCHD1 protein interactions provide an explanation for its mode of action on distinct loci regulated by SMCHD1, suggesting an important role for SUMOylation in the context of SMCHD1-mediated epigenetic regulation.

## Material and Methods

### Cell lines and culture

HeLa, HeLa1222, HEK293T, U2OS and hTERT-RPE1 cells were maintained in Dulbecco’s Modified Eagle Medium (DMEM)-Glutamax (Gibco 31966-047) supplemented with 10% heat inactivated Fetal Bovine Serum (FBS) (Gibco 10270-106) and 1% Penicillin-Streptomycin (Pen/Strep, Gibco 15140-122). HeLa1222 stably expresses His -SUMO2 and was described previously ^26^.

Primary and immortalized myoblast were maintained in Ham’s F10 Nutrient Mix-Glutamax (Gibco 41550-088) supplemented with 20% FBS, 1% Penstrep, 10ng/ml rhFGF (Promega G5071) and 1mM dexamethasone (Sigma-Aldrich D2915). For differentiation of myoblast cultures into myotubes, medium of confluent myoblast cultures was replaced with DMEM-Glutamax supplemented with 1% Pen/Strep and 2% KnockOut serum replacement (Gibco 10828-028) and cells were allowed to fuse for 48-72 hours.

Identity of HEK293T, HeLa and U2OS cells was validated and confirmed using in-house STR profiling. All cell lines used in the study were negative for mycoplasma infection as judged by regular testing (Mycoalert, Lonza LT07-318).

### Plasmids and antibodies

The lentiviral pRRL-CMV construct containing the full-length coding sequence for *SMCHD1* (pLV-*SMCHD1*) (described in ^25^) was used in this study. Addition of N-terminal tags in pLV-*SMCHD1* was done by addition of a multiple cloning site (MCS) by annealed oligo cloning, followed by insertion of either an HA-, 3xHA-, 3xFLAG, or 3xTY1-tag in the MCS. For introduction of missense variants at substrate lysine residues in SMCHD1, either full plasmid amplification site-direction mutagenesis using PfuUltra II Fusion HS DNA polymerase (Agilent) or overlap extension site-direction mutagenesis using Phusion DNA polymerase (NEB) was utilized. In either case, mutagenesis primer design was assisted by use of the PrimerX webtool (http://www.bioinformatics.org/primerx/). All clones were sequence verified by Sanger sequencing (Macrogen).

pLV-HA-*SENP1* and pLV-HA-*SENP2* were subcloned from coding sequence (CDS) clones (kind gift from Sabine Kuijpers, department of Cell and Chemical Biology – Leiden University Medical Center) to pRRL-CMV-HA backbones using *SacII* and *XbaI* restriction enzymes. For pLV-HA-*SENP5*, the full length *SENP5* transcript was amplified from HeLa cDNA using Phusion DNA polymerase (NEB) and cloned into the pRRL-CMV-HA backbone using *SacII* and *XbaI* restriction sites. The open reading frames of all plasmids were verified by Sanger sequencing.

The overexpression plasmid for GFP-TRIM28 was acquired from the Addgene plasmid repository (#65397)^27^.

Plasmids containing e.g. tagged SMCHD1 and SENP1, SENP2, and SENP5 are deposited in the Addgene plasmid repository.

Constructs encoding various shRNAs were obtained from the Mission shRNA library (TRC1, TRC1.5 and TRC2) (Sigma Aldrich) and are listed in table 2.

Primary antibodies used in this study are listed in table 3.

### Lentivirus production and transduction

Lentiviral particles were generated using a third-generation lentiviral system utilizing VSV-G as described previously ^28^. In brief, HEK293T cells were transfected with 3 packaging plasmids and the transfer plasmid encoding the CDS or shRNA of interest. 48h and 72h post transfection, medium was harvested, which was centrifuged at 2500RCF for 10 minutes and filtered through 45µm Acrodisc Tuffryn filters (Pall corporation). If necessary, harvested viral media was layered on a 20% Sucrose cushion and concentrated at 30.000 RPM for 2h in an Optima XE-100 ultracentrifuge (Beckman coulter). The HIV Type 1 p24 ELISA kit (Zeptometrix) was used for assessing the concentration of viral particles in all of the generated inoculates.

Prior to lentiviral transduction, cell medium was supplemented with positively-charged polycations in the form of DEAE-dextran (myoblasts) or polybrene (all other cell types used). Unless indicated otherwise, transductions were equalized to a viral load of 3ng viral particles per cm^2^ of culture surface, as determined by p24 ELISA. 16 hours post transduction, cell medium was refreshed, and cells were selected for puromycin (Sigma P8833) resistance for at least 48 hours.

### His-Pulldown of SUMO conjugates

His_10_ purification of SUMO conjugates was done according to Hendriks et. al ^26^ with minor modifications. Cells stably expressing His_10_-SUMO2 or His_10_-SUMO1 were washed and scraped in ice-cold PBS, and pelleted after a total lysate sample of 5% of input material was obtained. Cell pellets were rapidly lysed in lysis buffer (6M Guanadine-HCl, 100mM Na_2_HPO_4_/NaH_2_PO_4_, 10mM Tris-HCl, pH 8.0) and snap frozen in liquid nitrogen. Upon thawing, cell lysates were sonicated (3x 5s pulses, 30 Watt output power) using a microtip sonicator and lysates were normalized for protein concentration as measured by the Pierce BCA kit (Thermofisher 23225). Equalized lysates were then supplemented with β-mercaptoethanol (5mM) and imidazole (50mM) and incubated overnight with 20 µl (40 µl slurry) washed Ni-NTA beads (Qiagen). After washing with a wash buffer series (1x wash buffer 1: 6 M guanidine-HCl, 0.1M Na_2_HPO_4_/NaH_2_PO_4_ pH 8.0, 0.01M Tris-HCl pH 8.0, 10 mM imidazole pH 8.0, 5 mM β-mercaptoethanol, 0.1% Triton X-100. 1x wash buffer 2: 8 M urea, 0.1M Na_2_HPO_4_/NaH_2_PO_4_ pH 8.0, 0.01M Tris-HCl pH 8.0, 10 mM imidazole pH 8.0, 5 mM β-mercaptoethanol, 0.1% Triton X-100. 1x wash buffer 3: 8 M urea, 0.1M Na_2_HPO_4_/NaH_2_PO_4_ pH 6.3, 0.01M Tris-HCl pH 6.3, 10 mM imidazole pH 7.0, 5 mM β-mercaptoethanol, 0.1% Triton X-100. 2x wash buffer 4: 8 M urea, 0.1M Na_2_HPO_4_/NaH_2_PO_4_ pH 6.3, 0.01M Tris-HCl pH 6.3, 5 mM β-mercaptoethanol, 0.05% Triton X-100.), proteins were eluted from the beads by addition of 20 µl elution buffer (8M Urea, 100mM Na_2_HPO_4_/NaH_2_PO_4_, 500mM imidazole, 10mM Tris-HCl, pH7.0). Proteins were eluted 2x 30 minutes and lysates were pooled. For western blot analysis 10 µl 5x Laemmli sample buffer (10% SDS, 50% glycerol, 10% β-mercaptoethanol and 0.05% bromophenol blue in 300mM Tris pH6.8) was added.

### Western blot

For western blots of total cell lysates, cells were lysed in 1x Laemmli sample buffer (2% SDS, 10% glycerol, 60 mM Tris pH6.8) and equalized by Pierce BCA kit, after which lysates were supplemented with 2% β-mercaptoethanol and 0.01% bromophenol blue prior to boiling 10 minutes at 95°C. Depending on the desired separation pattern, samples were run on Criterion TGX or Tris-Acetate gels, or Mini-protean TGX gels (Bio-Rad), and transferred to Immobilon-FL PVDF membrane (EMB Millipore). Membranes were blocked in 4% skim milk (Sigma-Aldrich) and probed with primary antibody in 4% skim milk or Immunobooster (Takara). After washing, membranes were incubated with donkey-α-Rabbit IRDye-800CW, donkey-α-Mouse IRDye-680RD, Westernsure HRP goat-α-Mouse or Westernsure HRP goat-α-Rabbit (LI-COR). Immunocomplexes utilizing IRdye were visualized using an Odyssey Classic infrared imaging system (LI-COR) and analysed, and if quantified if necessary, using the accompanying application software (V3.0). HRP conjugated membranes were incubated in Pierce ECL Plus Western Blotting Substrate (Thermofisher 32132) and visualized by X-ray films, a C-Digit Blot Scanner (LI-COR) or an Amersham Imager 680 (GE Healthcare Life sciences). For antibodies used, we refer to table 3.

### Quantitative real-time PCR

Cells were lysed directly in QIAzol Lysis reagent (Qiagen 79306), and total RNA was extracted using the Direct-zol RNA miniprep kit (Zymo Research R2062), as per manufacturer’s instructions, including DNAseI treatment. RNA concentrations were determined by Nanodrop, and up to 2 µg RNA was used per sample for first strand cDNA synthesis using the RevertAid First Strand cDNA Synthesis Kit (K1622, ThermoFisher), as per manufacturer’s instructions. Synthesized cDNA samples in which *DUX4* expression was to be measured were treated with 2 units Ambion RNase H (ThermoFisher

AM2293) for 20 minutes at 37°C. Gene expression was measured using gene specific primers (See table 1), using iQ SYBR-Green Supermix (Bio-Rad) in a CFX-384 Real-Time PCR system (Bio-Rad). Data was analysed using the accompanying CFX-manager V3.1 software, followed by statistical analysis in GraphPad Prism 8.

### Microscopy

Cells were seeded on #1.5 glass slides (Hecht Assistant) or Screenstar 96 well plates coated with collagen-I. Cells were fixed for 10 minutes with 4% Paraformaldehyde (Electron Microscopy Sciences), and permeabilized with 0.3% Triton X-100 for 15 minutes. After blocking in PBS supplemented with 0.1% BSA and 0.05% Tween-20, primary antibodies were incubated for 1 hour at RT in blocking buffer. Slides were washed 3 times in PBS-T (0.05%) and incubated with secondary antibodies and DAPI (62248 ThermoFisher) for 1 hour at RT. Slides were washed 3 times in PBS-T (0.05%) and after airdrying, mounted on glass slides in Prolong Gold (P36930, ThermoFisher), while 96 well plates were stored with PBS. Image acquisition was performed using either a Leica SP8 upright confocal system, or a Leica SP8 White Light Laser (WLL) inverted confocal system (Leica). For antibodies used, refer to table 3. For automated high content image analysis, acquired images were analysed using a custom analysis pipeline in Cell Profiler software (version 4.2.5), using identical processing parameters for each image and treatment condition.

### Stability assays

HEK293T cells were reverse transfected in bulk using Polyethylenimine (PEI)(Polysciences), after which equal amount of cell suspension was seeded in 6-well culture plates. 24 hours post-transfection (PTF), cells were treated for the indicated times with 100µg/ml Cycloheximide (Sigma-Aldrich C4859) or 20µM MG132 (Sigma-Aldrich M7449), with DMSO as a vehicle control. After cell lysis in Laemmli lysis buffer, equal sample amounts were analysed by western blot as described.

### siRNA knockdown

For knockdown of gene expression by siRNA, cells were seeded into culture flasks and allowed to attach. After 24 hours, cells were transfected with siRNA (30pM/6 well) using Lipofectamine RNAiMAX Transfection Reagent (13778150, ThermoFisher). All siRNAs used in this study were acquired from Dharmacon. *SENP5*: siGENOME D-005946-02, D-005946-03; *SMCHD1*: siGENOME SMARTpool M-032684-00-0005; *T R I M*: *2*si*8*GENOME SMARTpool M-005046-01-0005; *H N R N*: *P K* siGENOME SMARTpool M-011692-00-0005; *SETDB1*: siGENOME SMARTpool M-020070-00-0005. An siRNA targeting luciferase (D-002050-01-20) was used as a negative control.

### Chromatin Immunoprecipitation (ChIP)

ChIP was performed as described earlier ^6,25^. In brief, for ChIP of endogenous proteins: cells were fixed directly in culture by supplementation of the media with formaldehyde to a 1% end concentration for 10 minutes. Fixation was quenched by addition of 1M Glycine to a final concentration of 125mM, and cells were washed in ice-cold PBS and collected. Nuclei were released by lysis in NP-Buffer, after which chromatin was sheared by use of a Bioruptor Pico (Diagenode) (15 cycles, 30 seconds on/off). Shearing efficiency of each sample was checked by phenol purification of sheared chromatin, and 60µg of pre-cleared chromatin was used per antibody. For each chromatin sample, an input sample was obtained representing 10% of IP volume. After overnight incubation of chromatin and antibodies, Protein A/G Sepharose beads (17-5280-01, 17-0618-01 - GE Healthcare (3:1 ratio A/G beads)) were added for 2 hours to precipitate antibody bound complexes. Beads were washed sequentially in low salt buffer, high salt buffer, lithium-chloride buffer and a double TE-buffer wash step. DNA was eluted by addition of 10% Chelex-100 (Bio-Rad) and denaturing of complexes by heating to 95°C for 10 minutes. ChIP/Input DNA samples were analysed using primer sets for specific genomic regions (Table 1) using iQ SYBR-Green Super mix (Bio-Rad) on a CFX384 Real-Time PCR system (Bio-Rad). ChIP samples were normalized to input by use of ΔCt calculations. Statistical analysis was performed in GraphPad Prism 7.

### Gene editing by CRISPR-Cas9

For generation of *SMCHD1* knock-out cell lines, the CRISPR-Cas9 system as published by Ran et. al. was utilized ^29^. A guide targeting exon 3 of *SMCHD1* (ACTGATTGACCGACTGTAGC) was designed with the help of the MIT CRISPR design tool (www.CRISPR.mit.edu), and cloned into the PX458 vector (Addgene #48138) ^29^. Cells were transiently transfected with the specific sgRNA plasmid or empty PX458 plasmid using PEI (HeLa) and sorted single-cell into 96 wells plate 48h PTF using a BD Aria III cell sorter (BD-Bioscience). Clones were screened by western blot for protein expression and Sanger sequencing for the mutation at the intended locus. Single cell clones of the empty PX458 transfection were sorted as clonal controls.

### Co-immunoprecipitation

Co-Immunoprecipitation was performed as described earlier ^30^. The method was modified depending on the nature of the target protein to be precipitated. For ectopically expressed constructs containing a GFP- or 3xHA-tag, HEK293T, U2OS or HeLa cells were transfected using polyethylenimine (PEI) (23966-2 Polysciences) at a 1:3 plasmid to PEI ratio. 48 hours post transfection, cells were washed twice with ice-cold PBS and lysed in NP40 lysis buffer (50 mM Tris-HCl pH 8.0, 150 mM NaCl, 1% Igepal CA-630) supplemented with cOmplete Protease Inhibitors, NaF and 20mM N-ethyl maleimide (NEM). Lysates were lysed while rotating at 4°C for 15 minutes, and were then centrifuged at maximum speed in a cooled microcentrifuge to pellet insoluble debris. An aliquot of lysate was taken as input and supplemented with 5x Laemmli Sample Buffer (10% SDS, 50% glycerol, 10% β-mercaptoethanol and 0.05% bromophenol blue in 300 mM Tris pH 6.8). The rest of the lysate was incubated with, depending on target tag, 8 µl washed HA-bead slurry (A2095-1ML, Sigma-Aldrich), 8 µl washed Flag M2 Affinity gel (A2220, Sigma-Aldrich), or 10 µl washed GFP-Trap Agarose beads (gta-20 Chromotek) for 2 hours while rotating on a tumbler at 4°C. Beads were washed 3x 5 minutes with lysis buffer, and subsequently boiled in 40 µl 2x Laemmli Sample buffer. Similar processing steps were used for endogenous Co-IP experiments. Antibodies were conjugated to a mixture of 20 µl Dynabeads protein A and protein G (3:1 ratio) (10002D and 10003D, Thermo Fisher) per sample. Antibodies (1 µg per Co-IP sample) were conjugated to washed Dynabeads for 1 hour prior to addition of cell lysates as indicated. At the same time, samples were precleared with a similar volume of A/G Dynabeads without addition of an antibody.

For co-immunoprecipitation of mass-spectrometry samples, we used one independent dish of transfected cells for a semi-biological triplicate of the Co-IP. Each set of MS samples consisted of 3 dishes with GFP as a negative control, and 3 dishes of GFP-SMCHD1 as target. Samples were processed as above, except that 20 µl washed GFP-Trap Agarose beads (gta-20 Chromotek). After washing with lysis buffer, beads were washed 2x with 50 mM ammonium bicarbonate and trypsinized overnight in 250 µl 50 mM Ammonium Bicarbonate containing 2.5 µg sequencing grade trypsin (V5111 Promega). Peptides were cleaned up by loading on in-house prepared tC18 stage-tips and washing with acetic acid solution. After elution with acetonitrile, samples were lyophilized and stored until analysis.

### Mass spectrometry data acquisition

Mass spectrometry data was acquired as in ^31^. In brief, Liquid Chromatography was performed on an EASY-nLC 1000 system (Proxeon, Odense, Denmark) connected to a Q-Exactive Orbitrap (Thermo Fisher Scientific, Germany) through a nano-electrospray ion source. The Q-Exactive was coupled to a 15 cm analytical column with an inner-diameter of 75 μm, in-house packed with 1.9 μm C18-AQ beads (Reprospher-DE, Pur, Dr. Maish, Ammerbuch-Entringen, Germany).

The chromatography gradient length was 100 minutes from 2% to 30% acetonitrile followed by 10 minutes to 95% acetonitrile in 0.1% formic acid at a flow rate of 200 nL/minute. The mass spectrometer was operated in data-dependent acquisition (DDA) mode with a top-7 method. Full-scan MS spectra were acquired in a range from 300 to 1600 m/z at a target value of 3 × 10^6^ and a resolution of 70,000, and the Higher-Collisional Dissociation (HCD) tandem mass spectra (MS/MS) were recorded at a target value of 1 × 10^5^ and with a resolution of 17,500 with a normalized collision energy (NCE) of 25%. The maximum MS1 and MS2 injection times were 250 ms and 120 ms, respectively. The precursor ion masses of scanned ions were dynamically excluded (DE) from MS/MS analysis for 40 sec. Ions with charge 1, and greater than 6 were excluded from triggering MS2 analysis.

### Mass spectrometry data analysis

To determine enrichment, LC-MS/MS Raw files were analyzed using MaxQuant software (v1.6.7.0) according to ^32^ using default settings. We performed the search against an in silico digested UniProt reference proteome for Homo sapiens (29th Aug 2022). Maximum number of mis-cleavages by trypsin/p was set to 3 and Carbamidomethyl(C) was deactivated as fixed modification. Oxidation (M) and Acetyl (Protein N-term) were left as variable modifications with a maximum number of 3. Label-Free Quantification (LFQ) was activated not enabling Fast LFQ. Minimum amount of peptides for quantification was set to 1. Match-between-runs feature was enabled with 0.7 min match time window and 20 min alignment time window.

MaxQuant proteingroups.txt file was further processed in the Perseus Computational Platform according to Tyanova et. al. ^33^. Values were log2 transformed, and potential contaminants and reverse peptides were removed. Samples were grouped in experimental categories and proteins not identified in every replicate in at least one condition were removed. Missing values were imputed using normally distributed values with 0.3 width and 1.8 down shift considering the total matrix. After imputation, the different conditions groups were statistically compared using two-tailed t-test with correction for multiple testing using permutation-based False Discovery Rate (FDR) equal to 0.05 with 200 permutations and an S0= 0.1. Results were exported to Microsoft Excel 365 (Microsoft, USA) for comprehensive visualization and browsing of the data.

### Mass spectrometry data availability

The mass spectrometry proteomics data have been deposited to the ProteomeXchange Consortium via the PRIDE partner repository ^34^ with the dataset identifier PXD055295.

## Results

### DUX4 expression in myogenic cells is regulated by SUMOylation

Previous studies have shown that SUMO regulates *Dux* expression in mouse ES cells and that hypoSUMOylation leads to derepression of *Dux* in a PRC1.6 and TRIM28/SETDB1 dependent manner^21^. We therefore asked if SUMO also regulates DUX4 in myogenic cells, the tissue showing misexpression of DUX4 in FSHD. We treated control and (genetically confirmed) FSHD1 and FSHD2 immortalized myoblasts with ML-792 ^35^, a small molecule inhibitor of SUMO activating enzyme (SAE) E1 ligase, and measured *DUX4* expression levels. The control myoblasts carry a 4qter allele of intermediate size (13 D4Z4 repeat units) with a somatic *DUX4* polyadenylation signal (i.e. a permissive FSHD allele), theoretically capable of *DUX4* expression in the context of SMCHD1 mutations ^36^, while the FSHD myoblasts are a diverse representation of genetic causes for FSHD. After ML-792 treatment *DUX4* (Figure 1A) and DUX4 target-gene (Supp. Figure 1A) expression was increased significantly in all myocytes. Interestingly, expression of the SMCHD1 regulated FSHD2 gene *LRIF1* was significantly upregulated by ML-792 treatment solely in the SMCHD1 missense FSHD2 myoblasts (Supp. Figure 1A). Because *DUX4* and DUX4 target gene expression increases with myogenic differentiation ^25^, myogenic markers *MYOG* and *MYH3* (Supp. Figure 1A) were measured. Differentiation was variably delayed in the presence of ML-792 reinforcing the role of SUMO on the regulation of DUX4. Similar to ML-792 treatment, inhibition of SUMOylation by depleting the SUMO E1 ligase component *UBA2* led to *DUX4* derepression in FSHD-patient derived myocytes (Supp Figure 1B). These results show that SUMOylation regulates *DUX4* expression in myogenic cells when the D4Z4 repeat chromatin structure is sensitized to *DUX4* expression, as is the case for FSHD and for unaffected carriers of a D4Z4 allele capable of expressing DUX4 in the context of FSHD2 mutations.

**Figure 1:**
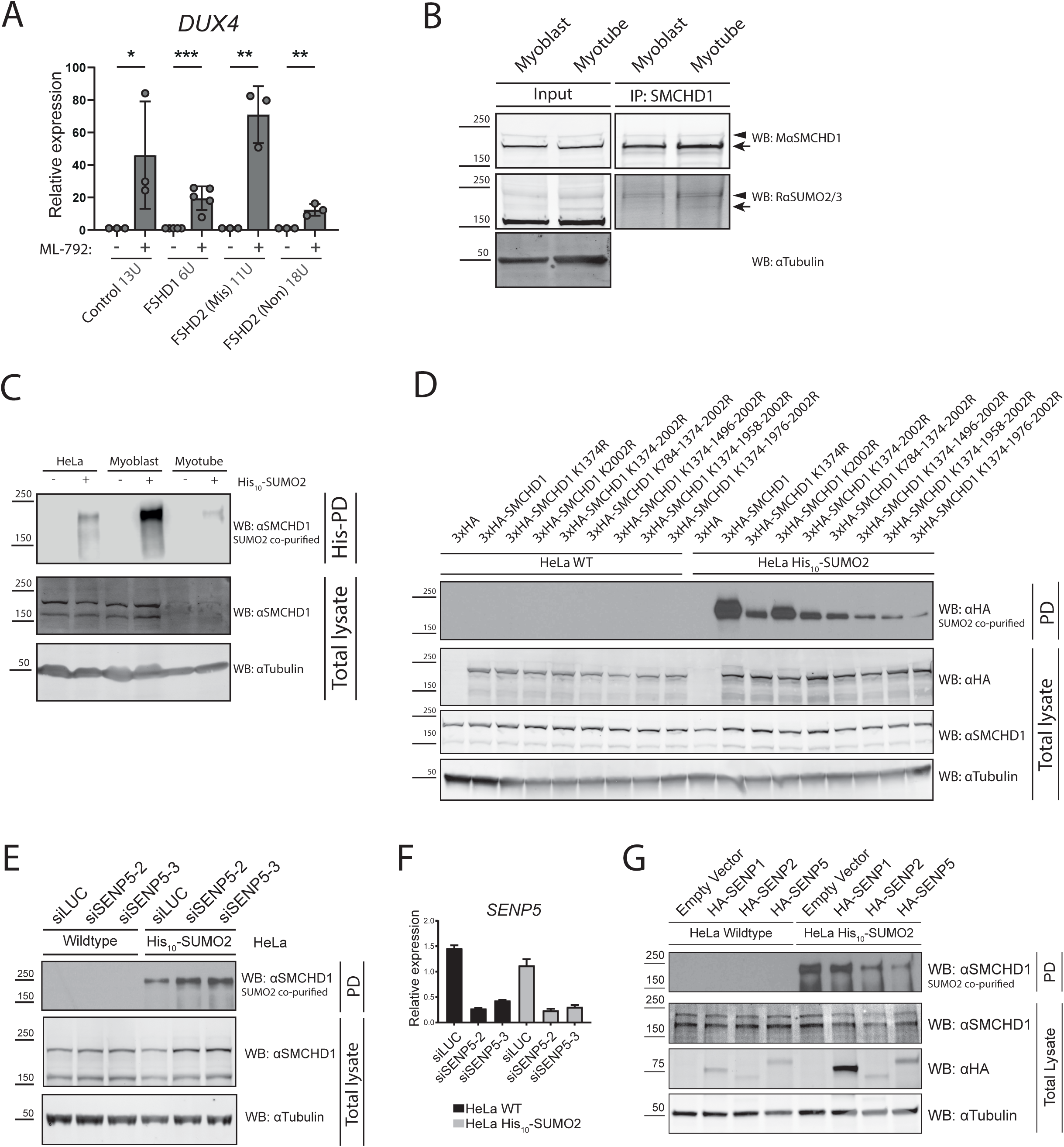
SMCHD1 is SUMOylated on lysine 1374 in different cell types. A: RT-qPCR analysis of *DUX4* expression in various FSHD-patient derived myotubes treated with ML-792. A control cell line carrying a permissive allele, as well as an FSHD1, FSHD2 (SMCHD1 missense mutation) and FSHD2 (SMCHD1 nonsense mutation) are used. (N=3) B: Immunoprecipitation analysis of endogenous SMCHD1 from control myoblast and myotube protein lysates. Antibodies specific for SMCHD1 and SUMO2/3 identify higher molecular weight bands (arrowheads) with overlapping SMCHD1 and SUMO2/3 signals. The arrow indicates the position of unmodified SMCHD1 protein. C: Western blot analysis of endogenous SMCHD1 enriched after His_10_-pulldown in HeLa cells, myoblasts and myotubes stably expressing His_10_-SUMO2, indicating the level of SMCHD1-SUMOylation. D: Western blot analysis of ectopically expressed 3xHA-SMCHD1 variants enriched after His_10_ purification (PD) of HeLa-(His_10_-SUMO2) cells, indicating the level of SMCHD1 SUMOylation. SMCHD1 variants include: K1374R, K2002R, K1374-2002R, K784-1374-2002R, K1374-1496-2002R, K1374-1958-2002R and K1374-1976-2002R. E: Western blot analysis of endogenous SMCHD1 enrichment in eluates after His_10_-SUMO2 pulldown in HeLa-(His_10_-SUMO2) *SENP5* depleted cells using specific siRNAs as indicated, siLUC was used as non-targeting control. F: RT-qPCR analysis of *SENP5* knockdown efficiency in siLUC and siSENP5 transfected HeLa wildtype and His_10_-SUMO2 cells used for His_10_ purification in Fig. 1E. G: Western blot analysis of endogenous SMCHD1 enrichment purified by His_10_-SUMO2 pulldown after overexpression of HA-tagged SENP1, SENP2 or SENP5 in HeLa-(His_10_-SUMO2) cells. RT-qPCR data was normalized to GUSB expression. Error bars: SEM. (*: P value <0.05, **: P value <0.01, ***: P value <0.001, ****: P value <0.0001, NS: Not-Significant – Students T-test). Tubulin serves as a loading control for all total protein lysates. PD: His_10_-pulldown lysate.

### SMCHD1 is SUMOylated in different cell types at multiple lysine residues

A global SUMOylome study in HeLa cells identified the SUMO2 modification of SMCHD1 ^5^, while in another study SMCHD1 was found modified by SUMO1 in the mouse brain ^3^. We hypothesized that transcriptional derepression of *DUX4* by ML-792 treatment or *UBA2* depletion might be a consequence of reduced SUMOylation of SMCHD1, or of its complex members, resulting in loss of the repressive function of SMCHD1. To investigate the SUMOylation of SMCHD1 specifically, we used HeLa cell lines stably expressing His_10_-SUMO1 or His_10_-SUMO2. After purification of SUMOylated proteins with nickel-NTA beads, western blot analysis of the pulldown (PD) lysates demonstrated the presence of endogenous SUMO1- and SUMO2-conjugated SMCHD1, indicating detectable levels of SMCHD1 SUMOylation in these cell lines (Supp. Figure 1C).

To test whether SMCHD1 can also be SUMOylated in myocytes we performed an SMCHD1 immunoprecipitation (IP) experiment in KM155 control immortalized myoblasts and myotubes ^37^. We observed that purified SMCHD1 shows a distinct signal for SUMO2/3 as a mass shifted band recognized by both SMCHD1 and SUMO2/3 antibodies (Figure 1B). This observation was confirmed by stably integrating His_10_-SUMO2-IRES-GFP into KM155 control myocytes and performing a His_10_- SUMO2 pulldown from proliferating and differentiated myocytes (Figure 1C). Immunoblotting for endogenous SMCHD1 confirmed the presence of SMCHD1-SUMO2 conjugates in myoblasts and myotubes (Figure 1C). Although basal SMCHD1 protein levels are reduced upon differentiation into myotubes (Figure 1C, input panel), which is in agreement with published data ^25^, SUMOylated SMCHD1 can still be detected.

We extended our SUMOylation analysis of SMCHD1 to different cell types. Using a cross-linked IP approach that preserves SUMO modifications during protein purification, we identified SMCHD1 SUMOylation by SUMO2/3 in HEK293T, HeLa, and U2OS cells providing experimental support that these cells can be used for mechanistic studies of SMCHD1 SUMOylation (Supp. Figure 1D).

Proteomics studies have provided evidence for SMCHD1 SUMOylation at six lysine residues: K784, K1374, K1496, K1958, K1976 and K2002 ^38^. To validate the SUMOylation sites of SMCHD1, we mutated single or multiple putative SUMO acceptor lysines to arginine in a 3xHA-SMCHD1 expression vector. The 3xHA tag contains no lysine residues minimizing the risk of SUMOylation artefacts. We next investigated the presence of SUMOylation on the putative SUMO acceptor lysines in SMCHD1 and observed a dramatic loss of SMCHD1 SUMOylation upon mutation of K1374 (Figure 1D), indicating that K1374 is the major SUMO2 acceptor site. Further substitutions of K1496, K1958 and K1976 resulted in further loss of SUMOylation of SMCHD1 (Figure 1D, right 3 lanes). Substitution of K784 and K2002 had no additional effect on SUMOylation of SMCHD1 by SUMO2 (Figure 1D).

Lysine residues can also be modified by ubiquitination (Ub). To rule out a potential role of ubiquitination at K1374 we used an HA-IP under denaturing conditions in HEK293T cells overexpressing either WT or K1374R mutant 3xHA-SMCHD1. Western blot analysis showed a shifted high-molecular weight band for wild-type, but not K1374R-mutant SMCHD1 (Supp. Figure 1E, arrowheads) which overlaps with a strong signal for SUMO2/3 probed on the same membrane. Probing for ubiquitinated proteins using the anti-ubiquitin FK2 antibody showed that both wild-type and K1374R SMCHD1 are equally modified by ubiquitin (Supp. Figure 1E). This indicates that while SMCHD1 can be both ubiquitinated and SUMOylated, K1374 is a specific substrate for SUMOylation. This observation is in accordance with published MS data which identifies ubiquitination on lysine K177, K240, K812 and K1256 of SMCHD1, although this has not been experimentally validated ^39^.

Finally, we investigated the direct effect of ML-792 on SMCHD1 SUMOylation and observed that an increasing concentration of ML-792 leads to a potent loss of SUMO2 on SMCHD1 without effecting total protein levels (Supp. Figure 1F).

Altogether we show that SMCHD1 is SUMOylated in different cell types and that SUMO2/3 modification of lysine 1374 of SMCHD1 is a conserved cellular process, maintained in all tested mammalian cell types.

### SUMO protease SENP5 modulates SMCHD1 SUMOylation

Sentrin/SUMO specific proteases (SENPs) are responsible for activating newly produced SUMO moieties and removing SUMO from acceptor proteins ^40^. There are six different SENPs (SENP1/2/3/5/6/7) that are ubiquitously expressed. To gain more insight into the molecular mechanism of SMCHD1 SUMOylation, we individually knocked down each SENP by lentiviral transduction of shRNA in HeLa cells and tested the effect on SMCHD1 SUMOylation. We found that only knockdown of *SENP5* resulted in a robust increase of SMCHD1 SUMOylation, as shown by the enhanced signal after His_10_-SUMO2 pulldown (Supp. Figure 2A), while knockdown of other SENPs had no reproducible effect on SMCHD1 SUMOylation (data not shown). Knockdown of *SENP5* was validated by RT-qPCR (Supp. Figure 2B), as no reliable antibody for western blot analysis was available. To confirm the effect of *SENP5* knock down, we independently knocked down *SENP5* in HeLa cells using two individual siRNAs and observed an increase of SUMO2-SMCHD1 after SUMO2-His_10_ purification (Figure 1E and F). Finally, we overexpressed HA-tagged SENP1, SENP2, and SENP5 in HeLa cells and found a decrease in SMCHD1 SUMOylation upon overexpression of HA-SENP2 and HA-SENP5, but not HA-SENP1 (Figure 1G). Overexpression of SENP5 almost completely abolished SMCHD1 SUMOylation while overexpression of SENP2 had an intermediate effect.

SENP5 is predominantly localised in the nucleolus ^41^, while SMCHD1 is localized in the nucleoplasm, enriched at the Xi territory in female cells but largely excluded from nucleoli ^15^. To gain insight into the detailed subcellular localization of SMCHD1, we employed Stimulated Emission Depletion (STED) super-resolution IF microscopy of endogenous SMCHD1 in U2OS cells and observed bright SMCHD1 foci in the nucleolar periphery (Supp. Figure 2C). IF staining of ectopically expressed HA-SENP5 in U2OS cells confirmed its localization in nucleoli, together with a diffuse nucleoplasmic signal (Supp. Figure 2D) ^42^.

Finally, to investigate the effect of *SENP5* knockdown on DUX4, we depleted *SENP5* using siRNA in FSHD1 and FSHD2 myoblasts and observed a significant downregulation of *DUX4* and DUX4 target gene expression (Supp. Figure 3A and 3B), again indicating that modifying SUMOylation (by SENP5) affects the repression of the D4Z4 repeat. Depletion of *SENP5* in a control myoblast line incapable of *DUX4* expression due to absence of the somatic *DUX4* polyadenylation signal, showed no consistent effect on *DUX4* or DUX4 target genes (Supp. Figure 3C), indicating that the changes in expression of these targets in the FSHD cell lines is DUX4-dependent.

Altogether our data suggests that SENP5 and SMCHD1 colocalize in the nucleolar periphery, that primarily SENP5 activity modulates SMCHD1 SUMOylation, and that SENP5 can modulate *DUX4* expression in FSHD myogenic cells, likely through its effect on SMCHD1 SUMOylation.

### SUMOylation of SMCHD1 does not affect the protein’s stability, chromatin binding or dimerization properties

SUMOylation can increase protein stability through protection from degradation by the proteasome or decrease stability by recruitment of SUMO-targeting ubiquitin ligases (STUbL) ^43,44^. Therefore, we studied whether SUMOylation protects SMCHD1 from degradation by the proteasome. We introduced wild-type (WT) or K1374R mutant SMCHD1 into SMCHD1-KO HEK293T cells by lentiviral transduction. Expression levels of WT or mutant SMCHD1 in the rescued SMCHD1-KO HEK293T cells (referred to as WT Rescue and K1374R Rescue from hereon) were comparable to endogenous SMCHD1 levels (Supp. Figure 4A). WT Rescue or K1374R Rescue cells were cultured in the presence of the proteasomal inhibitor MG132 and harvested at different time points. We observed that after 24 hours of treatment both wild-type and K1374R mutant SMCHD1 protein accumulated in similar quantities as detected by semi-quantitative western blot (Figure 2A and 2B), indicating that SUMOylation of SMCHD1 at K1374 does not affect protein stability by attenuation of proteasomal degradation in this experimental setup.

**Figure 2:**
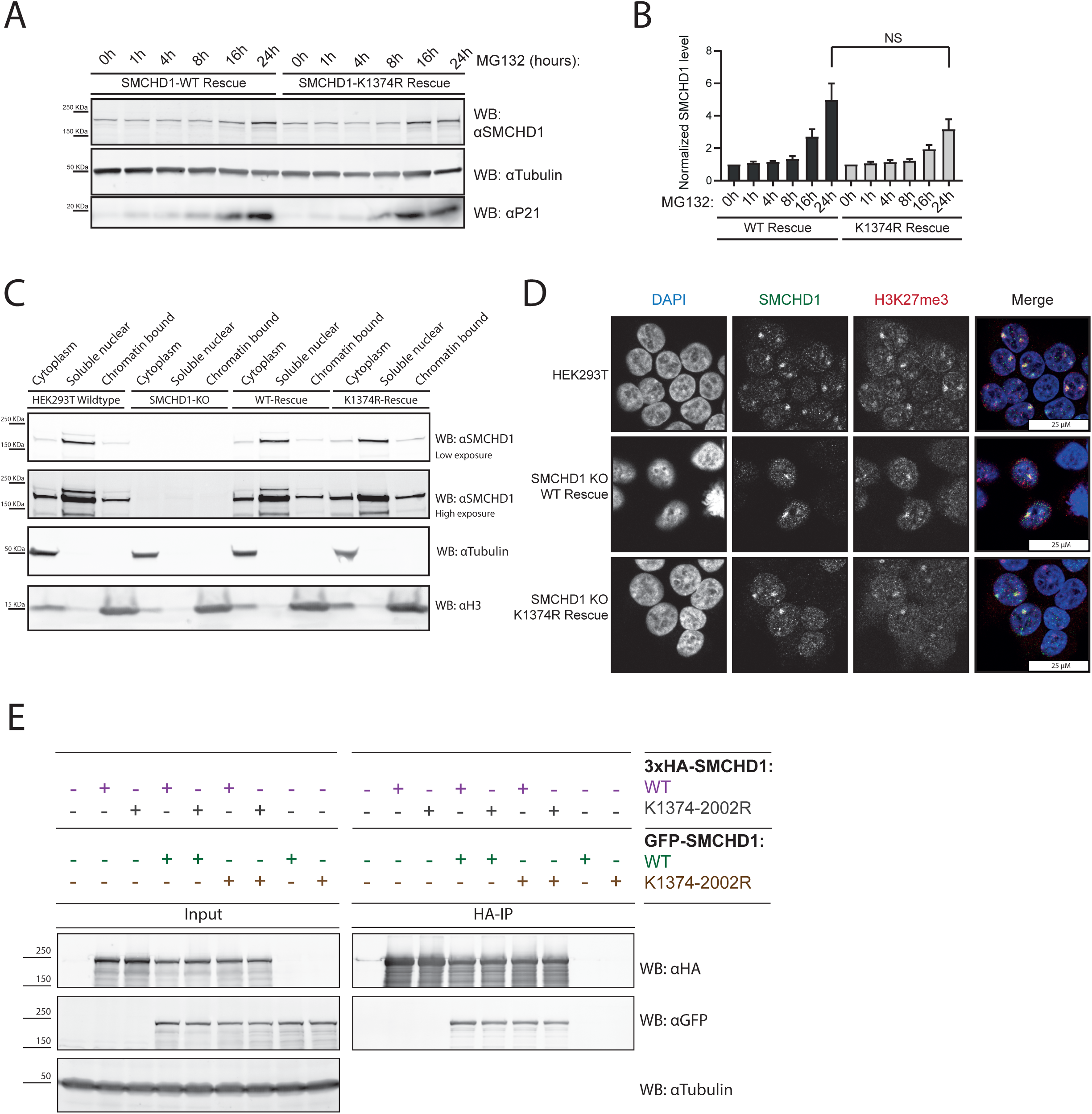
deSUMOylation does not affect SMCHD1 protein characteristics. A: Western blot analysis of SMCHD1 protein levels in MG132 treated HEK293T SMCHD1-KO cell lines rescued with stably expressed SMCHD1-WT and SMCHD1-K1374R . Protein P21 was probed as a positive control for the MG132 treatment efficacy. B: Quantification of the western blot in panel A, with SMCHD1 signal normalized to tubulin. C: Western blot analysis of cellular fractionations of HEK293T wildtype, SMCHD1-KO and rescue lines as indicated. Tubulin and H3 are used to respectively demonstrate purity of the cytoplasmic and chromatin fractions. D: Immunofluorescent confocal microscopy analysis of SMCHD1 enrichment at the Xi in HEK293T SMCHD1-KO rescue lines. H3K27me3 staining was used as a marker for Xi localization, DAPI counterstains the nucleus. Scalebar: 25 µm. E: Western blot analysis of GFP-SMCHD1 co-precipitating with 3xHA-SMCHD1 after purification of HA-tagged proteins. 3xHA- or GFP-tagged SMCHD1 WT or K1374-2002R variants were overexpressed as indicated. Tubulin serves as a loading control of input lysates. Error bars: SEM. (*: P value <0.05, **: P value <0.01, ***: P value <0.001, ****: P value <0.0001, NS: Not-Significant – One-Way ANOVA)

To further test whether the stability of SMCHD1 is affected by mutating lysine 1374, WT Rescue and K1374R Rescue cells were treated with cycloheximide (CHX) to inhibit translation, and cells were harvested at different time points. Quantification of SMCHD1 protein levels showed again no significant difference between wild-type and K1374R SMCHD1 over time (Supp. Figure 4B and 4C). Similarly, HEK293T cells transiently expressing 3xHA-SMCHD1 WT, K1374R, K2002R and K1374-2002R and treated with CHX showed no difference in protein stability between the SMCHD1 variants after 24 hours of treatment (Supp. Figure 4D and 4E). Altogether we conclude that SUMOylation at K1374 does not have a major role in SMCHD1 protein stability.

SMCHD1 is a chromatin binding protein. To investigate if SMCHD1 SUMOylation regulates global chromatin association we compared the chromatin-bound and soluble nuclear fraction of WT and K1374R mutant SMCHD1 after subcellular fractionation. WT SMCHD1 and K1374R SMCHD1 showed similar distributions in these fractions (Figure 2C), indicating that these SMCHD1 mutants are not defective in their subnuclear distribution. SMCHD1 was shown by several studies to associate wthe inactive X (Xi) ^15^. To investigate whether SUMOylation of SMCHD1 affects its recruitment to Xi chromatin, we analysed the enrichment of SMCHD1 at the Xi (Barr body), which can be visualised as bright nuclear H3K27me3 foci ^23^. In WT HEK293T cells, these bright H3K27me3 foci colocalized with SMCHD1 similar to the WT and K1374 rescue cell lines (Figure 2D), indicating that SUMOylation of K1374 alone is not essential for SMCHD1 recruitment to Xi.

Dimerization of SMCHD1 through its hinge domain is essential for chromatin association ^22^. We therefore asked whether impaired SUMOylation indirectly influences the hetero- or homodimerization of SMCHD1. HEK293T cells were transiently transfected with GFP or 3xHA-tagged WT or K1374-2002R SMCHD1 in different combinations. After immunoprecipitation of 3xHA-SMCHD1 variants using HA-agarose beads, interacting GFP-SMCHD1 WT or K1374-2002R variants were detected in all combinations of WT and K1374R-K2002R (Figure 2E), indicating that the SMCHD1 variant lacking its primary SUMOylation site was able to form both hetero- and homodimers similar to WT SMCHD1. To test whether chromatin association by SMCHD1 dimers was affected after the loss of the primary SUMOylation site, we performed ChIP-qPCR analysis of known SMCHD1-binding loci in WT Rescue and K1374R Rescue SMCHD1 KO cells. SMCHD1 binding at the five interrogated genomic loci (SMCHD1 binding sites *D4Z4-DR1*, *DUX4-Q*, *GAPDH, LRIF1* promoter and *WWC3*) was comparable in WT Rescue and K1374R Rescue cells (Supp. Figure 4F). These observations suggest that loss of SUMOylation at K1374 does not impair the dimerization or chromatin association of SMCHD1.

Altogether, we found no evidence that loss of SUMOylation of SMCHD1 K1374 alters its stability, chromatin binding or self-dimerization properties although it remains possible that subtle changes in any of these parameters cannot be captured in our experimental design.

### Identification of SUMO dependent interactome of SMCHD1

SUMOylation also regulates a wide range of cellular processes by modulating protein-protein interactions e.g. through SUMO interaction motifs (SIMs) found in proteins ^2^. Several studies have identified SMCHD1 interacting proteins ^22,30^, but none of these studies used cysteine protease inhibitors, inadvertently losing the reversible SUMO modification during the procedure. N-Ethylmaleimide (NEM) is a cysteine protease inhibitor that effectively preserves SUMOylation by inhibiting the catalytic activity of SENPs. To identify the SUMO-dependent interactome of SMCHD1 we therefore performed mass spectrometry (MS) analysis of immunoprecipitated GFP-SMCHD1 obtained under SUMO preserving lysis conditions in different cell types. We compared the interactome of three independent purifications of GFP-SMCHD1 and GFP as a control in HEK293T, HeLa and U2OS cell lines, which yielded respectively 882, 690 and 480 significantly enriched SMCHD1 interaction partners. Of these, 323 interactions co-occurred in all three cell lines (Figure 3A-C, Suppl. figure 5A and B and Supplementary table 4).

**Figure 3:**
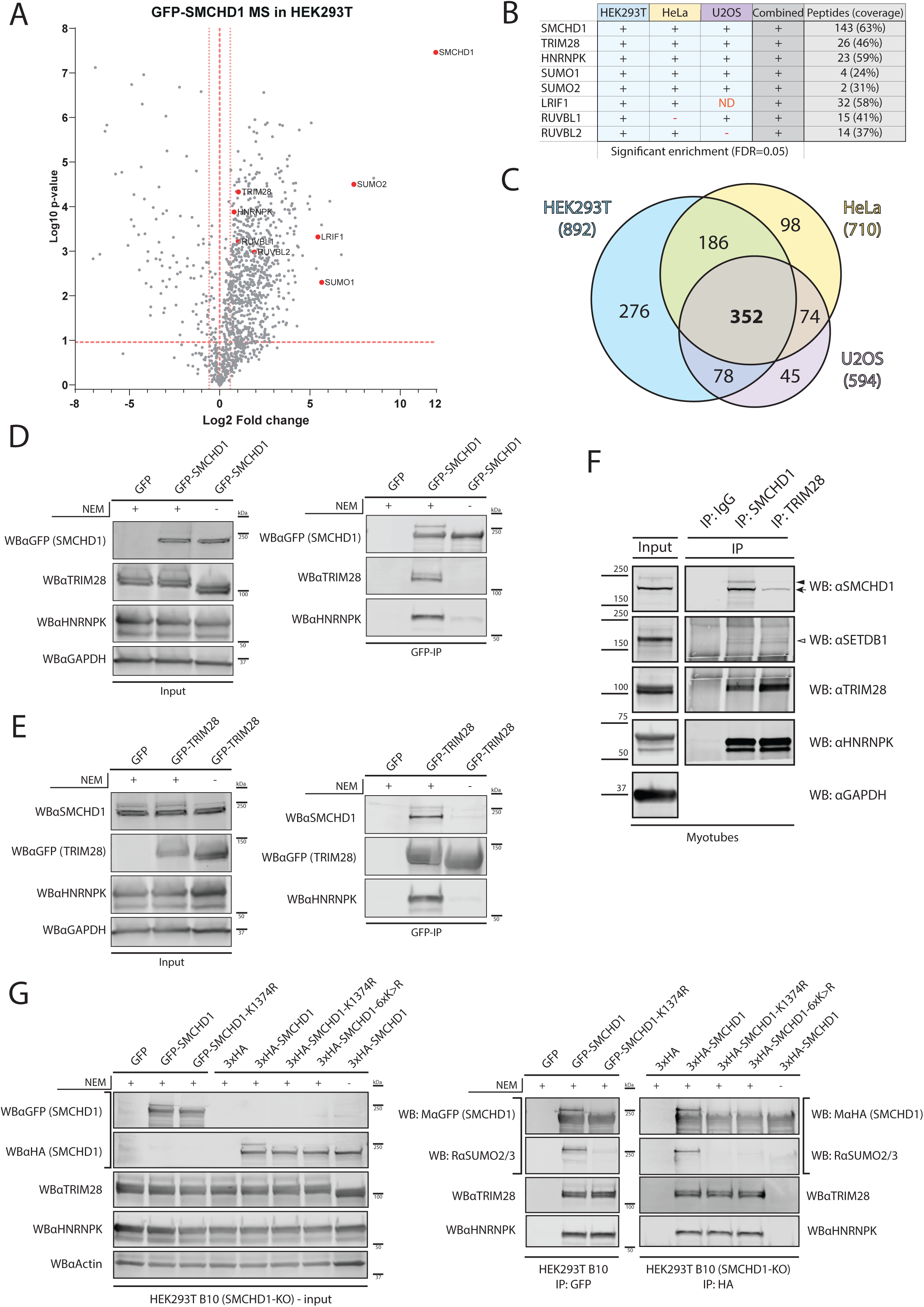
Identification of a SUMOylation dependent complex between SMCHD1, TRIM28, HNRNPK and SETDB1. A: Volcano plot visualizing the SMCHD1-interacting proteome of GFP-SMCHD1 vs GFP in transfected HEK293T cells analysed using LC-MS (N=3), using SUMO-preserving lysis conditions. Horizontal dashed line indicates the Log10 P-value significance treshold. Vertical dotted line indicates 1,5x fold change. Proteins of interest are highlighted in red. B: Summary table of mass spectrometry data visualized in (A), representing the pulldown of GFP vs. GFP-SMCHD1 expressing HEK293T cells (N=3). Identical experiments were simultaneously performed in HeLa (N=3) and U2OS cells (N=3). Table shows whether the proteins of interest (TRIM28, HNRNPK, SUMO1 and SUMO2) and known interactors of SMCHD1 (LRIF1, RUVBL1, RUVBL2) were detected with significant confidence over the respective control in each cell line used. LRIF1 is not expressed in U2OS. C: Venn diagram of mass spectrometry data acquired from HEK293T, HeLa and U2OS. Numbers between brackets indicate the total number of significantly enriched SMCHD1 interacting proteins per cell line. Numbers inside the Venn diagram indicate overlap of enriched protein, as indicated by the overlap between the circles. D: Western blot analysis of GFP-SMCHD1 interacting proteins under SUMO-preserving conditions (+NEM) compared to SUMO-lossy conditions (-NEM) in GFP vs. GFP-SMCHD1 transfected HEK293T cells. Left panel: whole cell lysate input. Right panel: enrichment after GFP-immunoprecipitation (GFP-IP). E: Western blot analysis of SMCHD1-interacting proteins using a reciprocal immunoprecipitation of GFP vs. GFP-TRIM28 in transfected HEK293T cells under SUMO-preserving (+NEM) and SUMO-lossy (-NEM) conditions. Left panel: input, right panel: GFP-IP. F: Western blot analysis of SMCHD1-interacting proteins in myotubes after native SMCHD1 and native TRIM28 immunoprecipitation in SUMO-preserving conditions (+NEM). Solid arrowhead: SUMOylated SMCHD1. Arrow: SMCHD1. Open arrowhead: SETDB1. IgG: binding specificity control. G: Western blot analysis of interacting proteins of GFP-SMCHD1(WT) compared to GFP-SMCHD1-K1374R mutant, as well asHA-SMCHD1(WT), HA-SMCHD1-6xK mutant after GFP- and HA-IP as indicated. HEK293T SMCHD1 knock out cells (HEK293T clone B10) were lysed under SUMO-preserving (+NEM) and SUMO-lossy (-NEM) conditions. Actin is used as a loading control. LC-MS: liquid chromatography mass spectrometry. NEM: N-Ethylmaleimide. ND: Not detected. FDR: False Discovery Rate. GAPDH is used as a loading control.

Besides known interaction partners of SMCHD1, like LRIF1 ^23,45^, RUVBL1 and RUVBL2 ^30^, we identified TRIM28 and HNRNPK in the SUMO-protected interactome of SMCHD1 in all three cell lines (Figure 3A, 3B). Both proteins are known to localize to SMCHD1 target loci but have not been identified as SMCHD1 protein interactors. TRIM28 is a component of D4Z4 chromatin and represses DUX4 in somatic cells ^46^ while HNRNPK facilitates SMCHD1 recruitment to the Xi through binding to Xist ^47,48^ supporting a biological function of their interaction. We confirmed the MS data in reciprocal co-immunoprecipitation experiments with HEK293T cells transiently expressing GFP-SMCHD1 or GFP-TRIM28 under SUMO preserving conditions. These results show that TRIM28 is present in the GFP-SMCHD1 IP fraction (figure 3D) and SMCHD1 can be detected in the GFP-TRIM28 IP fraction (figure 3E) while co-precipitated HNRNPK is detected in both the GFP-TRIM28 and GFP-SMCHD1 IP samples (Figure 3D, E). SMCHD1 interactions with TRIM28 and HNRNPK were lost when SUMOylation was not preserved (-NEM samples, Figure 3D, E). The SUMO-dependent SMCHD1 interaction with TRIM28 and HNRNPK was also detected in immortalized myoblast cell lines by endogenous co-immunoprecipitation (Figure 3F). Interestingly SETDB1, a histone methyltransferase and co-repressor of TRIM28, although not significantly enriched in the SMCHD1-interactome dataset, was also found in the SUMO-preserved IP fractions (Figure 3F), confirming an earlier study reporting this interaction ^49^.

Next, we studied whether the SMCHD1-TRIM28-HNRNPK interaction depends on the SUMOylation of SMCHD1 exclusively. We transiently expressed GFP-tagged and 3xHA-tagged versions of wild type SMCHD1 and SUMO-defective K1374R SMCHD1 in HEK293T SMCHD1 KO cells. Additionally, we expressed the 3xHA-tagged SMCHD1-6xK>R variant, in which the six most prominent SUMO acceptor lysines (K784, K1374, K1496, K1958, K1976 and K2002) are replaced by arginine ^38^. We analysed GFP- and HA-immunoprecipitated fractions by western blot. One HA-IP-sample was prepared without the addition of NEM to serve as a control for loss of SUMOylation. TRIM28 and HNRNPK were detected in all IP fractions prepared under SUMO-preserving conditions, suggesting that while their mutual interaction is SUMO-dependent, this interaction does not critically depend on SMCHD1 SUMOylation alone (Figure 3G).

TRIM28 is a known SUMO E3 ligase ^50^. Considering the interaction of SMCHD1 and TRIM28, we speculated that TRIM28 might have a direct role in SMCHD1 SUMOylation. We therefore knocked down TRIM28 in our His_10_-SUMO2 expressing HeLa cell lines using shRNA and performed a His_10_- purification of SUMOylated protein. We observed a reduction in SUMOylated SMCHD1 upon knockdown of TRIM28 (Figure 4A). Reciprocally, upon ectopic expression of GFP-TRIM28 in the same cell line, we could purify a noticeably larger amount of SUMOylated SMCHD1 (Figure 4B). Together, this data indicates that SMCHD1 interacts with and gets SUMOylated by TRIM28.

**Figure 4:**
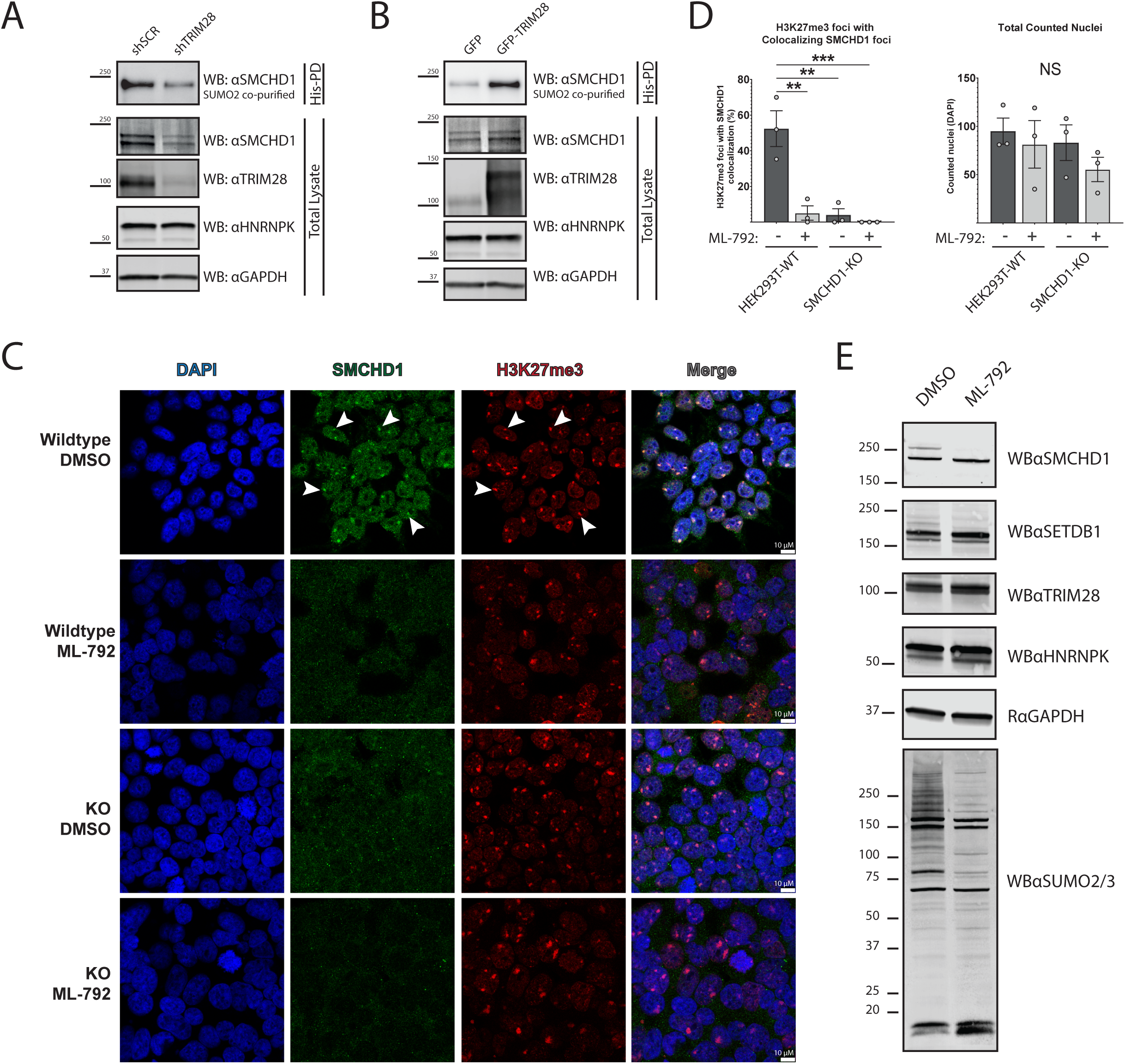
TRIM28 regulates SMCHD1 SUMOylation, and SUMOylation is essential for SMCHD1 localization to Xi. A: Western blot analysis of endogenous SMCHD1 enrichment in eluates after His_10_-SUMO2 pulldown in HeLa-(His_10_-SUMO2) cells knocked-down for *TRIM28* using siRNAs as indicated, siLUC was used as non-targeting control. B: Western blot analysis of endogenous SMCHD1 enrichment purified by His_10_-SUMO2 pulldown after overexpression of GFP-tagged TRIM28 in HeLa-(His_10_-SUMO2) cells. C: Immunofluorescent microscopy of SMCHD1 enrichment at the Xi in HEK293T SMCHD1 wildtype and SMCHD1-KO cell lines, treated with ML-792 or DMSO for 48 hours as indicated. H3K27me3 staining is a marker for Xi localization, DAPI counterstains the nucleus. Scalebar: 25 µm. N=3 D: Quantification of ML-792 or DMSO treated HEK293T SMCHD1 wildtype and SMCHD1-KO IF images shown in C. H3K27me3 foci were scored for colocalization of SMCHD1 foci. The percentage of H3K27me3 with a corresponding SMCHD1 signal is shown in the panel, while the number of counted nuclei is shown on the right. Each dot represents the quantification of a set of IF images from independent replicates. Error bars: SEM. (**: P value <0.01, ***: P value <0.001, NS: Not-Significant – One-Way ANOVA). E: Western blot analysis of SMCHD1-interacting protein complex components in total lysates of HEK293T wildtype cells treated for 48 hours with ML-792 as in B.

### SMCHD1 localization to Xi is SUMO-dependent

A relationship between HNRNPK function and SMCHD1 recruitment to Xi has been suggested previously ^48^ in mouse embryonic fibroblasts, where it was observed that knockdown of Hnrnpk resulted in the loss of SMCHD1 at Xi. Direct recruitment of SMCHD1 by HNRNPK was not considered, as the authors did not find an interaction between these proteins in the absence of SUMO-preserving conditions. To investigate whether SMCHD1 localization to Xi in humans is dependent on any of the SUMO-dependent interaction partners, we depleted SMCHD1, TRIM28, HNRNPK and SETDB1 in hTERT-RPE1 and HEK293T cells with siRNAs and analysed SMCHD1 recruitment to the Xi using immunofluorescence microscopy, where Xi was visualized as bright H3K27me3-positive foci (Supp. Figure 6A and B). Except for the positive control (siSMCHD1), only knockdown of HNRNPK resulted in a reduction of SMCHD1 localization to Xi foci (Supp. Figure 6A and B), confirming previous observations ^48^. We next treated HEK293T cells with SUMOylation inhibitor ML-792 and assessed SMCHD1 localization to Xi (Figure 4C-D and Supp. Figure S7). Strikingly, while ML-792 treatment had no effect on the protein levels of SMCHD1, SETDB1, TRIM28 and HNRNPK (Figure 4E), inhibiting SUMOylation almost fully abolished SMCHD1 localization at the Xi, indicating that a major defect in SMCHD1 recruitment or maintenance at the Xi occurs upon loss of SUMOylation (Figure 4C-D). The major defect in Xi territory integrity is further exemplified by the remarkable reduction in H3K27me3 foci in ML-792 treated cells, indicating a broader role for SUMOylation in the integrity of Xi territories.

Altogether we found that TRIM28, HNRNPK, SETDB1 and SMCHD1 interact with each other in a SUMO-dependent fashion and that this interaction is not solely dependent on the SUMOylation of SMCHD1. However, SUMOylation is critical for the Xi chromatin structure, including the presence of SMCHD1 at the inactive X chromosome.

### SUMOylation regulates gene expression of SMCHD1 target loci differently in myocytes compared to HEK293T cells

In our previous experiments we observed that ML-792 mediated depletion of SUMOylated proteins resulted in various degrees of *DUX4* gene dysregulation in FSHD1 myocytes. We therefore optimized the delivery of the SUMO-inhibiting agent ML-792 in FSHD1 myocytes to assess the gene expression response of selected loci in more detail. We noticed a potent response on total SUMO protein conjugation in an FSHD1 cell line induced by the entire dose range of ML-792 (Supp. Figure 8A), as well as a clear dose-dependent increase of *DUX4* and DUX4 target genes *ZSCAN4* and *KHDC1L* expression (Figure 5A and Supp. Figure 8C). This loss of *DUX4* repression by ML-792 was not caused by loss of SMCHD1 protein (Supp. Figure 8A), nor was it caused by an increased induction of myogenic differentiation, a known driver of *DUX4* expression in FSHD myocytes (Supp. Figure 8C – *MYOG* and *MYH3* panels). ML-792 treatment did not lead to an increase of SMCHD1-regulated single copy loci *LRIF1* (Figure 5A) or *WWC3* (Supp figure 8C) expression at any treatment dose, while an increase in *PCDHB2,* a member of the protocadherin gene cluster locus known to be regulated by SMCHD1 ^13^, was only observed at higher ML-792 doses (Supp Figure 8C)).

**Figure 5:**
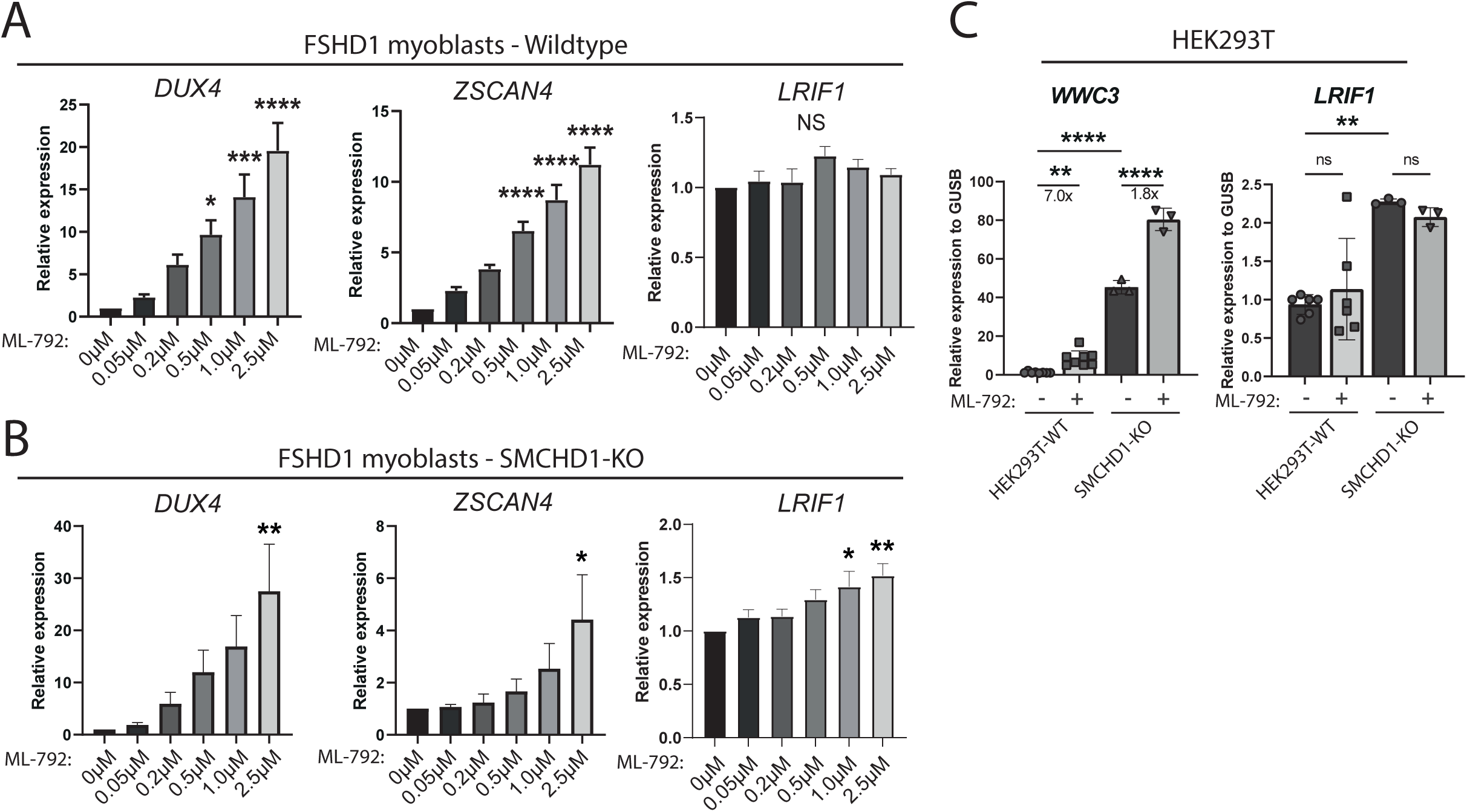
The SUMO-dependent interaction partners of SMCHD1 repress various loci of the genome. A: RT-qPCR analysis of *DUX4, ZSCAN4* and *LRIF1* expression in FSHD1 patient-derived myoblasts treated with SUMOylation inhibitor ML-792 at indicated concentrations. (N=5) B: RT-qPCR analysis of *DUX4, ZSCAN4* and *LRIF1* expression in FSHD1 patient-derived myoblasts knocked out for SMCHD1 treated with ML-792 at indicated concentrations. Two KO clones were independently treated 3 times. (N=6) C: RT-qPCR analysis of *WWC3* and*LRIF1* expression in HEK293T and HEK293T SMCHD1 KO cells treated with DMSO or ML-792 (WT: N=7, KO: N=3) (*: P value <0.05, **: P value <0.01, ***: P value <0.001, ****: P value <0.0001, NS: Not-Significant – Kruskall-wallis (A) and One-Way ANOVA)

To assess whether ML-792’s effect on SMCHD1 SUMOylation is a critical factor driving the upregulation of these genes, we also applied the ML-792 titration on two independent SMCHD1-KO myocytes (Supp. Figure 8B) derived from the same FSHD1 patient. *DUX4* and DUX4 target gene expression was still dose-dependently affected by ML-792 treatment as in SMCHD1 expressing FSHD1 myocytes (Figure 5B and Supp. Figure 8D), suggesting a non-exclusive role for SMCHD1 SUMOylation on DUX4 repression. Interestingly, we noticed a modest increase in *LRIF1* expression in response to ML-792 treatment in the SMCHD1-KO cells, which was absent in the SMCHD1-expressing myocytes (Figure 5A and B, *LRIF1* panel). This response in *LRIF1* expression is consistent with the effect observed in FSHD2 myocytes carrying an SMCHD1 missense mutation (Supp. Figure 1A). Treatment of HEK293T SMCHD1-WT and SMCHD1-KO cells with ML-792 did not show any SMCHD1-dependent differences in expression of *PCDHB2, WWC3* and *LRIF1* (Figure 5C, Supp. Figure 8E), although the amplitude of the increased expression of *PCDHB2* and *WWC3* by ML-792 was dampened in cells lacking SMCHD1.

These observations suggest regulation of gene expression by SUMOylation differs in different cell types and on different loci within the same cell type, with a various degree of dependence on the presence of SMCHD1.

### The SMCHD1-TRIM28-HNRNPK-SETDB1 chromatin repressor complex is recruited via different mechanisms to single copy and repeat loci

Previously we studied the role of SMCHD1 as a repressor of *LRIF1* and *DUX4* as representative examples of autosomal single copy and repeat loci, respectively ^19^. We therefore investigated whether the recruitment of SMCHD1 to these loci depends on the presence of our identified binding partners or of the SUMOylation thereof. We first used specific siRNA pools to monitor the immediate response to depletion of SMCHD1, TRIM28, HNRNPK and SETDB1 on protein level (Figure 6A). An SMCHD1-KO HEK293T line was included to confirm antibody specificity for the ChIP-qPCR procedure. Semi-quantitative analysis of protein levels after siRNA knockdown revealed that none of the specific knockdowns changed the protein levels of any of the other complex partners (Supp. Figure 9A). Surprisingly, *LRIF1* expression was reduced in siHNRNPK treated cells whereas we confirmed the increased levels of *LRIF 1*upon loss of SMCHD1 (Figure 6B)^19^. Also, we observed a significant derepression of *WWC3* in SMCHD1 depleted cells (Supp. figure 9B). ChIP-qPCR experiments for SMCHD1 in HEK293T cells depleted for each of the interaction partners by siRNAs showed that *DUX4 Q* binding of SMCHD1 to *DUX4 Q* was not dependent on TRIM28, HNRNPK or SETDB1 (Figure 6C). However, on the *LRIF1* promoter, HNRNPK depletion resulted in significantly increased recruitment of SMCHD1, explaining the earlier observation of decreased *LRIF1* gene expression. As HNRNPK depletion showed a loss of SMCHD1 at the Xi territory, we also assessed the occupancy of SMCHD1 at the X-chromosomal *WWC3* locus. Here, a non-significant trend for SMCHD1 loss could be observed after HNRNPK depletion (Figure 6C).

**Figure 6:**
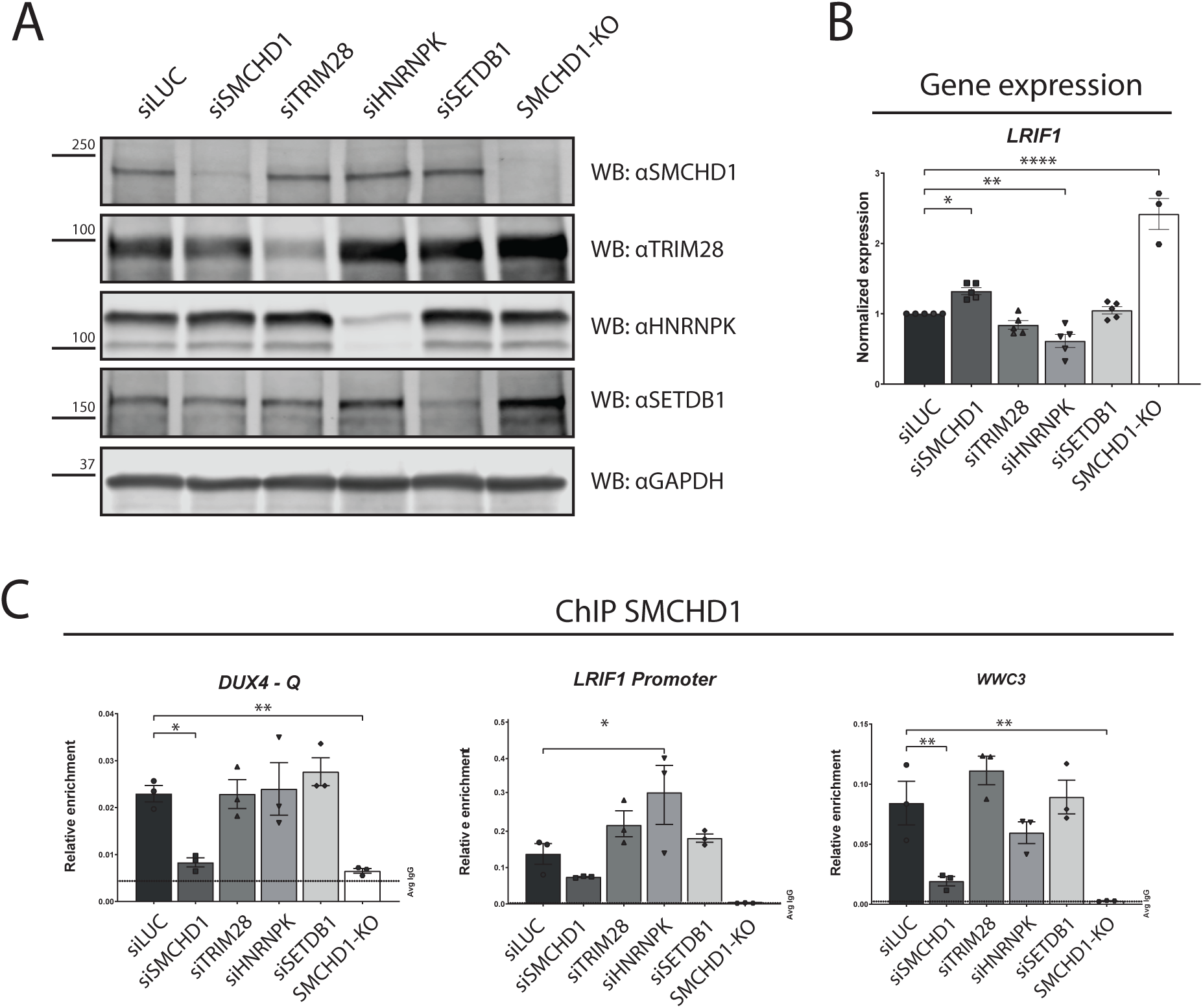
Chromatin association of SMCHD1 is differentially regulated at various loci with a partial dependence on its SUMO-dependent interaction partners. A: Western blot analysis of total lysates of HEK293T cells transfected with siRNAs targeting the indicated genes. SMCHD1-KO cells were not transfected with siRNA. GAPDH was used as a loading control. B: RT-qPCR analysis of *LRIF1* expression in HEK293T corresponding to the siRNA transfected samples in A. C: ChIP-qPCR analysis of SMCHD1 enrichment at the *D4Z4* (Q) repeat, *LRIF1* Promoter, and *WWC3* locus. Samples were depleted for gene expression by siRNA transfection as indicated and verified in A. Values are plotted as relative enrichment over input. IgG: negative control. Error bars: SEM. (*: P value <0.05, **: P value <0.01, ***: P value <0.001, ****: P value <0.0001, NS: Not-Significant – One-Way ANOVA).

Treatment of FSHD myocytes with SUMO-inhibitor ML-792 significantly increased expression of *DUX4* (Figure 1A, 5A-B). However, we did not observe any change in the enrichment of SMCHD1 bound to the interrogated D4Z4 loci, nor of TRIM28 or SETDB1 (Supp. Figure 10A). We observed, however, a 3-fold increase of SMCHD1 occupancy at the *LRIF1* promoter in ML-792 treated FSHD1 myocytes (Supp. Figure 10A). In HEK293T cells, modulation of SUMOylation by ML-792 treatment did not affect *LRIF1* expression (Figure 5C). This can indicate that while SUMO inhibits recruitment of SMCHD1 to the *LRIF1* promoter region, unSUMOylated SMCHD1 is impeded in properly regulating the activity of the locus. Similarly, a knockdown of SENP5 by siRNAs, which increases SMCHD1 SUMOylation, resulted in a loss of SMCHD1 on the LRIF1 promoter in HeLa cells, but had no effect on other loci measured (Supp. Figure 10B).

## Discussion

While the function of the chromatin repressor SMCHD1 has been the topic of several studies, the regulation of SMCHD1 levels and activity itself has remained enigmatic. Here, we report the ubiquitous post translational modification of SMCHD1 by SUMO. This study provides experimental support for several SUMO proteomics studies, all suggesting that SMCHD1 is among the most abundantly SUMOylated proteins ^3,38,51^. SUMO2/3 SUMOylation of SMCHD1 occurs predominantly at K1374, with a visible loss of higher molecular weight SMCHD1 upon its substitution. K1374 lies between the ATPase-Transducer domain and the hinge domain ^11^, a region of unknown functionality. We did not detect significant effects of SUMOylation on SMCHD1 protein stability, cellular localization or chromatin binding, but we do show that SUMOylation is important for its association with the Xi, for SMCHD1 protein interactions and that SUMOylation affects gene expression of loci controlled by SMCHD1. The fact that we observe residual SMCHD1 SUMOylation after mutating six different lysine residues can be explained by published mass spectrometry data suggesting that apart from the 6 major lysines we studied, SMCHD1 can be SUMOylated at another 15 minor SUMO acceptor lysines ^38^.

SMCHD1 is deSUMOylated by the SUMO-specific protease SENP5, as shown in reciprocal SENP5 knockdown and overexpression experiments. In support, regulation of SMCHD1 SUMOylation by SENP5 was recently also suggested by Dönig et al. in a siSENP5 proteomics approach ^52^. SENP5 is a nucleolar protein and our study provides evidence that SMCHD1 localizes in yet uncharacterized foci in the nucleolar periphery. Interestingly, the D4Z4 repeat localizes in nucleolar associated domains (NADs) as well as the lamina associated domains (LADs) ^53–55^, which are known sites of heterochromatin maintenance. Possibly, the SENP5-mediated deSUMOylation of D4Z4-bound SMCHD1 or its SUMO-dependent interacting proteins is concentrated in specific foci in the nucleolar periphery. Our data indicates that SMCHD1 SUMOylation is promoted by the E3 ligase activity of TRIM28, with a reciprocal response of SMCHD1 SUMOylation levels after depletion or ectopic expression of TRIM28 in HeLa cells. Furthermore, a recent study by Salas-Lloret *et al*. suggests that SMCHD1 is also processed by the PIAS4 E3 ligase ^56^, but this was not further experimentally validated.

### SUMOylation has a role in D4Z4 heterochromatin maintenance in somatic cells

We also report the importance of SUMOylation in silencing of *DUX4* in somatic cells. Treatment of cells with the SAE inhibitor ML-792 leads to a rapid, global loss of SUMOylated conjugates, and has been reported to upregulate *Dux* in mES cells ^21^. Alike *Dux,* we observed a dose-dependent upregulation of *DUX4* in myoblasts upon treatment with ML-792. *DUX4* derepression was reproduced by the knockdown of the SUMO E1 ligase *UBA2*, confirming the importance of SUMOylation in the maintenance of a repressive D4Z4 chromatin state in somatic cells.

While SMCHD1 is rapidly deSUMOylated after treatment with ML-792, SUMO-reduced SMCHD1 K1374R is not actively released from D4Z4 chromatin. As SMCHD1 is part of a SUMOylated complex of repressor proteins, it is likely that the loss of one member or the loss of SUMOylation of a single complex member does not completely disrupt the complex^2,57^. However, upon global loss of SUMOylation by ML-792 treatment, repressor complex activity at D4Z4 decreases and transcriptional silencing cannot be maintained. Reciprocally, impairment of active deSUMOylation by SENP5 knockdown leads to increased D4Z4 repression. This suggests that while SUMOylated SMCHD1 is of importance to the maintenance of the repressive D4Z4 chromatin state, its activity can be partially compensated, which is known to occur for SUMO-mediated protein complexes ^1^. It is also important to emphasize that while ML-792 treatment leads to a rapid loss of SUMOylation, this loss is the result of a disturbance in steady state SUMOylation levels caused by rapid cycles of SUMOylation and deSUMOylation. Intriguingly, ML-792 treatment leads to increased levels of SMCHD1 at the *LRIF1* promoter, a major site of SMCHD1 accumulation in the genome ^19^. The observation that SMCHD1 occupancy at the *LRIF1* promoter is decreased when knocking down SENP5 indicates that this locus might be more directly regulated by SUMOylation than the other loci we investigated.

### The SUMO-dependent SMCHD1-HNRNPK interaction

We identified novel interaction partners of SMCHD1 by mass spectrometry analysis after SUMO-preserving SMCHD1 enrichment. We were able to confirm several of these interactors (TRIM28 and HNRNPK, as well as their associated protein SETDB1 ^58^) by co-immunoprecipitation, and demonstrated that absence of NEM resulted in complete abolishment of SMCHD1 SUMOylation and protein interaction. HNRNPK is one of the more notable interaction partners that we validated as Jansz et. al. have previously found a relationship between HNRNPK activity and SMCHD1 recruitment to the Xi mediated by an interaction between HNRNPK and the lncRNA Xist ^48^. However, a direct interaction between SMCHD1 and HNRNPK was not demonstrated probably due to the absence of SUMO-preserving chemicals ^48^. Their suggested mechanism of SMCHD1 functionality at Xi through PRC1, H2AK119Ub and Xist may need to be revisited, as it can be simplified by the SUMO-dependent interaction between SMCHD1 and HNRNPK. Indeed, we demonstrated that SMCHD1 localization to the Xi is lost upon ML-792 treatment likely due to a failure of SMCHD1 recruitment by Xist-HNRNPK. ML-792 treatment causes large perturbations in the repressive histone marks seen on the Xi territory and we confirm that in human cells, alike murine cells ^48^, a dramatic loss of SMCHD1 occurs on the Xi after HNRNPK knockdown.

HNRNPK is a nuclear DNA and RNA binding protein with a wide variety of roles, such as enhancing or repressing gene transcription and regulation of lncRNAs ^59^. The reported binding of HNRNPK to Xist is also a suggested mechanism for SMCHD1’s activity to compact the higher order chromatin of the Xi ^16,18,47,48^. However, experimental limitations prevented us from addressing the function of HNRNPK at the D4Z4 repeat, and whether it is mediated through lncRNAs as well.

### The context-dependent function of the SUMO-mediated SMCHD1-interacting chromatin repressor complex

Our ChIP and expression data support a versatile model for SMCHD1-mediated repression through its interaction with TRIM28, HNRNPK and SETDB1 depending on the genomic context. All partners have been reported to be SUMOylated ^38^, and a functional relationship between HNRNPK and TRIM28 has been established ^58^. In mouse ES cells, SUMOylation levels of SMCHD1, TRIM28, DNMT3B, and members of the PRC1.6, PRC2 and NuRD complexes are high compared to somatic cells ^60^. SUMOylation of these repressive chromatin factors is suggested to regulate the transition between pluripotent and somatic cells, in part by maintaining SUMO-dependent *Dux* repression to prevent 2C activation ^21 60^. There is remarkable overlap of these SUMOylated pluripotency regulating genes and known FSHD genes ^6,61^, or D4Z4 regulatory proteins ^25,62^. This could suggest that these mechanisms are regulated similarly or are involved in establishing repressive chromatin at D4Z4 in early embryonic development.

At *autosomal repetitive loci* such as D4Z4, the data presented here shows strong evidence for the importance of SUMOylation in maintaining *DUX4* repression in somatic cells. Loss of SUMOylation in myoblasts has a profound and rapid effect on *DUX4* expression, which seems to be partially mediated by SMCHD1 deSUMOylation. The dramatic increase in *DUX4* levels upon treatment with ML-792 is reminiscent of the increase of *DUX4* seen upon transient knockdown of *SMCHD1* in myogenic cells derived from FSHD patients ^25^. This sensitivity of the D4Z4 repeat to loss of SUMO-based repression could help to explain the burst-like nature of DUX4 expression considering the dynamic nature of SUMOylation (Figure 7, top panel) ^36^.

**Figure 7:**
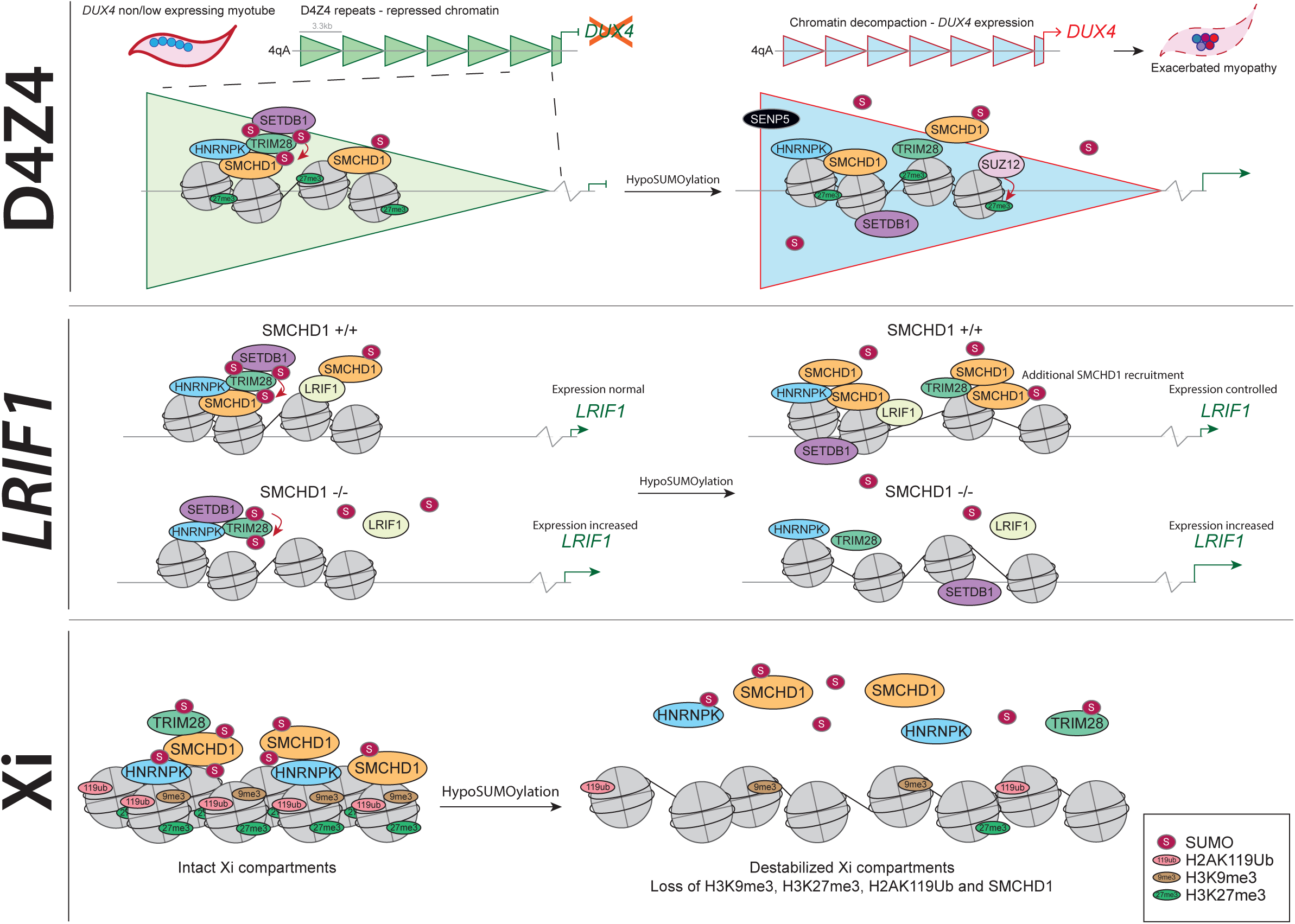
model of the role of SUMOylation in SMCHD1 function at various types of genomic loci. SMCHD1 is SUMOylated on K1374 by SUMO2/3, and interacts with chromatin repressor proteins HNRNPK, TRIM28 and SETDB1 in a SUMO dependent manner. The repetitive D4Z4 heterochromatin is maintained by a set of repressor complexes, of which several are known SUMO targets. HypoSUMOylation by cellular stressors of various nature can disrupt the homeostasis of the D4Z4-bound repressor complexes. Ultimately this leads to D4Z4 chromatin relaxation and inappropriate expression of DUX4 protein in skeletal muscle cells. The *LRIF1* locus is regulated by SMCHD1 binding its promoter and inhibiting transcription. Upon loss of SUMOylation of the SMCHD1 interacting protein repressor complex, additional SMCHD1 is recruited to *LRIF1* to maintain transcriptional control. In cells lacking SMCHD1 this compensation mechanism fails, leading to increased *LRIF1* expression. In a normal situation in female cells, the inactive X (Xi) chromosome is coated by repressive histone marks, *Xist*, and chromatin modifiers such as SMCHD1. Loss of SUMOylation leads to a defect in establishment or maintenance of proper Xi homeostasis and a loss of repressive components from the Barr body.

At *autosomal single copy loci* such as the *LRIF1* promoter SUMOylation also plays a role in expression regulation^45^. We recently showed that the *LRIF1* promoter is regulated by the interdependent binding of LRIF1 and SMCHD1 at this locus ^19^, with loss of SMCHD1 leading to high levels of *LRIF1* transcription. We now show that this interplay is layered with a SUMO regulatory function (Figure 7, middle panel), with a loss of SUMOylation triggering a reorganization of *LRIF1* chromatin. In SMCHD1 expressing myocytes this loss of SUMOylation at the *LRIF1* promoter seems to be partially compensated by additional recruitment of SMCHD1. However, in SMCHD1 haploinsufficient FSHD, or SMCHD1 knockout myocytes this compensatory mechanism is impaired resulting in increased *LRIF1* expression. The intricacies of the balance of LRIF1 recruitment on D4Z4 expression could add another layer of complexity to the SUMO-dependent regulation of *DUX4* repression.

At the *inactive X chromosome*we provide a novel mechanism for the previously described HNRNPK-dependent SMCHD1 recruitment to Xi and uncover a previously unconsidered role for SUMOylation in X inactivation and/or maintenance (Figure 7, lower panel). Our data shows that SMCHD1 and HNRNPK are members of a protein complex which is stabilized by SUMOylation, and that disruption of SUMOylation leads to the loss of SUMO-dependent SMCHD1-HNRNPK interaction and a dramatic reorganization of the chromatin structure of the Xi.

Altogether, our data suggests SUMO-dependent SMCHD1-mediated repression is organized differently at the three genomic loci – Xi, D4Z4 and *LRIF1* – and uncovers a previously undescribed role of SUMOylation in the establishment or maintenance of X inactivation.

## Acknowledgements

We would like to thank all patients and their families who contributed to this study. This study was supported by grants from the US National Institute of Arthritis and Musculoskeletal and Skin Diseases (R01AR066248) and the Prinses Beatrix Spierfonds (W.OR14-04; W.OR15–26). RG, MST, JB and SMvdM are members of the European Reference Network for Rare Neuromuscular Diseases [ERN EURO-NMD]. ACOV is the recipient of the European Research Council Grant (ERC; grant 310913). RG-P is recipient of a Young Investigator Grant from the Dutch Cancer Foundation (KWF-YIG 11367/2017-2) and a EMERGIA2020 grant from the Andalusian Government (EMRGIA20_00276). The authors acknowledge the Flow cytometry Core Facility (FCF) staff of Leiden University Medical Center (LUMC) in Leiden, the Netherlands (https://www.lumc.nl/research/facilities/fcf) for cell sorting assistance. Authors acknowledge the staff of the LUMC light microscopy facility for assistance with (confocal) microscopy experiments.

## Author contributions

RG, MST, JB and SMvdM designed and conceptualized the study. RG, MST and IMW performed experiments and processed data. KRS and PJvV performed supporting experiments and cloned full length *SMCHD1* plasmids. ACOV and RG-P assisted with generation and analysis of proteomics data. RG, MST, RG-P, ACOV, SJT, JB and SMvdM contributed to interpreting results. RG and MST drafted the manuscript. RG, MST and SMvdM finalized the manuscript. All authors read and approved the final manuscript.

## Competing interests statement

SMvdM is a Board member of Renogenyx and has acted as consultant and/or is a member of the advisory board for several pharmaceutical companies developing therapeutics for FSHD. SMvdM is co-inventor of several FSHD-related patent applications.

## Data availability statement

The data supporting the findings of this study are included in the main and supplementary figures. Proteomics data has been deposited in the PRIDE partner repository (PXD055295). Raw data can be supplied upon reasonable request to the corresponding author.

---

**Figure S1:** A: Additional RT-qPCR analysis of *DUX4* expression in various FSHD patient-derived myotubes treated with ML-792. A control cell line carrying a permissive allele, as well as an FSHD1, FSHD2 (SMCHD1 missense mutation) and FSHD2 (SMCHD1 nonsense mutation) are used. (N=3)

B: RT-PCR analysis of *DUX4* expression in myocytes depleted for *UBA2*. shSCR was used as non-targeting control (shControl). C: Western blot analysis of endogenous SMCHD1 after His_10_ purification of HeLa cell lines stably expressing His_10_-SUMO2 or His_10_-SUMO1. Covalently linked SMCHD1 is detected with αSMCHD1 antibody.

D: Western blot analysis of 3xHA-SMCHD1 purified from HEK293T, HeLa and U2OS under formaldehyde crosslinked conditions. Bottom panel (HA-IP) shows the overlapping signals for the SMCHD1 and SUMO2/3 antibody in the indicated cell lines.

E: Western blot analysis after HA-immunoprecipitation under denaturing conditions in HEK293T cell lines overexpressing 3xHA-SMCHD1 wildtype or K1374R mutant as indicated. Membrane was probed for SUMO2/3, total ubiquitin (Ubi-FK2) and HA-tag as indicated. Arrowheads indicate molecular weight-shifted SUMO-modified SMCHD1. Arrows mark unSUMOylated SMCHD1.

F: Western blot analysis of SUMOylated SMCHD1 purified by His_10_-SUMO2 pulldown after treatment of cells with various concentration of ML-792 as indicated.

Tubulin was used as loading control for western blots. RT-qPCR data was normalized to *GUSB* expression. Error bars: SEM. (*: P value <0.05, **: P value <0.01, ***: P value <0.001, ****: P value <0.0001, NS: Not-Significant – One-Way ANOVA (panel A) or Student T-test (panel B).

**Figure S2:** A: Western blot analysis of endogenous SMCHD1 enrichment in eluates after His_10_-SUMO2 pulldown in HeLa-(His_10_-SUMO2) cells knocked-down for *SENP5* using shRNA as indicated, shSCR was used as non-targeting control.

B: RT-qPCR analysis of *SENP5*expression of samples used in panel C confirms knockdown of *SENP5* by shRNAs targeting SENP5.

C: Stimulated Emission Depletion (STED) super resolution immunofluorescent analysis of SMCHD1 subcellular localization in the nucleus of U2OS cells. Foci of SMCHD1 localizing to the nucleolar periphery can be observed. Scalebar: 5 µm.

D: Confocal microscopy analysis of cells overexpressing HA-SENP5 confirms its correct localization to the nucleoli. No localization of HA-SENP5 to the inactive X counterstained by H3K27me3 was observed. Scalebar: 25 µm.

Tubulin was used as loading control for western blots. PD: His_10_-pulldown lysate. RT-qPCR data was normalized to *GUSB* expression.

**Figure S3:** A: RT-qPCR analysis of the expression of various transcripts in FSHD1 patient-derived myotubes (MB073) transfected with *SENP5* targeting siRNAs, or siLUC as control.

A: RT-qPCR analysis of the expression of various transcripts in FSHD2 patient-derived myotubes (MB200) transfected with *SENP5*targeting siRNAs, or siLUC as control.

A: RT-qPCR analysis of the expression of various transcripts in myotubes derived from a healthy control individual (MB135) transfected with *SENP5* targeting siRNAs, or siLUC as control.

RT-qPCR data was normalized to *GUSB* expression. Error bars: SEM. ND: Not detected. (*: P value

<0.05, **: P value <0.01, ***: P value <0.001, ****: P value <0.0001, NS: Not-Significant – One-Way ANOVA)

**Figure S4:** A: Western blot analysis of SMCHD1 expression in wildtype HEK293T, SMCHD1-KO, and SMCHD1 WT and K1374R rescue lines confirms KO of endogenous SMCHD1 and re-expression of integrated SMCHD1 constructs respectively.

B: HEK293T SMCHD1-WT or -K1374R rescue cells were treated with inhibitor of translation cycloheximide (CHX) as indicated and analysed by western blot for SMCHD1 levels. Doublets could be observed for wildtype but not the SMCHD1-K1374R substitution. Protein P21 was probed as a positive control for the CHX treatment efficacy.

C: Quantification of the western blot in panel A, with SMCHD1 signal normalized to tubulin.

D: HEK293T overexpressing 3xHA-tagged SMCHD1 or K1374R, K2002R and K1374-2002R treated with inhibitor of translation cycloheximide (CHX) as indicated and analysed by western blot for HA-SMCHD1 levels. Doublets could be observed for wildtype and K2002R, but not for mutants harbouring a K1374 substitution.

E: Quantification of the western blot presented in panel D. The 3xHA-SMCHD1 signal was normalized to actin and the ratio between 24h CHX (+) and 24h DMSO (-) was plotted.

F: ChIP-qPCR analysis of SMCHD1 enrichment at the D4Z4-DR1, D4Z4 – DUX4-Q, GAPDH and WWC3 loci in the HEK293T SMCHD1-KO and rescue lines as indicated. N = 2.

Arrows mark unSUMOylated SMCHD1, arrowheads mark molecular weight-shifted SUMOylated SMCHD1. Tubulin and Actin were used as loading control for western blots. Error bars: SEM. (NS: Not-Significant – One-Way ANOVA).

**Figure S5:** A, B, C: Volcano plots visualizing the SMCHD1-interacting proteome analysed using LC-MS of GFP-pulldown from whole-cell protein lysates of GFP vs. GFP-SMCHD1 transfected U2OS cells (A), HeLa cells (B) and a combined analysis of HEK293T, U2OS and HeLa cells (C) under SUMO-preserving conditions (N=3 for all). Horizontal dashed line indicates the threshold of Log10 P-value significance. Vertical dotted line indicates 1,5-fold change. Proteins of interest are highlighted in red. Of note: LRIF1 is not expressed in U2OS cells, therefore absent in (A).

**Figure S6:** Immunofluorescent confocal microscopy analysis of SMCHD1 enrichment at the Xi in hTERT-RPE1 (A) and HEK293T (B) cells treated with siRNAs targeting various genes as indicated. H3K27me3 staining was used as a marker for Xi localization, DAPI counterstains the nucleus. Scalebar: A: 10 µM, B: 50 µM.

**Figure S7:** A: Quantification of ML-792 or DMSO treated HEK293T SMCHD1 wildtype and SMCHD1-KO IF images shown in figure 4C. Left panel: Number of H3K27me3 foci colocalizing with SMCHD1 foci. Middle panel: Number of H3K27me3 foci not colocalizing with SMCHD1 foci. Right panel: Total number of H3K27me3 foci. Each dot represents the quantification of a set of IF images from independent replicates. (*: P value <0.05, ***: P value <0.001, NS: Not-Significant – One-Way ANOVA).

B: Summary data of all quantified data shown in figures 4C and S7A.

**Figure S8:** A: Western blot analysis of SUMO2/3 conjugates in FSHD1 myoblasts treated with SUMO inhibitor ML-792 to confirm treatment efficacy.

B: Western blot analysis of SMCHD1 expression in MB073 wildtype and 2 independent clones generated by CRISPR-Cas9, confirming knockout of SMCHD1 expression.

C: RT-qPCR analysis of expression of various indicated transcripts in FSHD1 patient-derived myoblasts treated with ML-792 at indicated concentrations. (N=5)

D: RT-qPCR analysis of expression of various indicated transcripts in FSHD1 patient-derived myoblasts knocked out for SMCHD1 treated with ML-792 at indicated concentrations. (N=6, 2 independent clones measured in technical triplicates).

E: RT-qPCR analysis of expression of *PCDHB2* in HEK293T WT and SMCHD1-KO cells treated with ML-792 as indicated.

RT-qPCR data was normalized to *GUSB* expression. (*: P value <0.05, **: P value <0.01, ***: P value

<0.001, ****: P value <0.0001, NS: Not-Significant – One-Way ANOVA)

**Figure S9:** A: Quantified normalized protein levels of the representative western blot shown in Figure 6A. Respective protein levels were normalized to housekeeping protein GAPDH.

B: RT-qPCR analysis of *SMCHD1, TRIM28, HNRNPK, SETDB1, PCDHB2* and *WWC3* expression in HEK293T cells after transfection with indicated siRNAs.

C: ChIP-qPCR analysis of SMCHD1 enrichment at *GAPDH* and *DUX4-DR1* region in HEK293T cells transfected with siRNAs targeting the indicated genes. SMCHD1-KO HEK293T cells were not transfected with siRNAs and serve as an experimental control. Values are plotted as relative enrichment over input. IgG values are plotted as a dashed line.

Error bars: SEM. ND: Not detected. (*: P value <0.05, **: P value <0.01, ***: P value <0.001, ****: P value <0.0001, NS: Not-Significant – One-Way ANOVA).

**Figure S10:** A: ChIP-qPCR analysis of SMCHD1 enrichment at *D4Z4* (Q and DR1) and the *LRIF1* promoter region, in FSHD1 patient-derived myoblasts treated with ML-792 at the indicated concentrations. Values are plotted as relative enrichment over input. Averaged IgG values are plotted as a dashed line. (N=2)

B: ChIP-qPCR analysis of SMCHD1 enrichment at *D4Z4* (Q), *GAPDH* and the *LRIF1* promoter region, in HeLa cells treated with siRNAs targeting SENP5 as indicated. Values are plotted as relative enrichment over input. Averaged IgG values are plotted as a dashed line. (N=1)

Error bars: SEM. ND: Not detected. (*: P value <0.05, **: P value <0.01, ***: P value <0.001, ****: P value <0.0001, NS: Not-Significant – One-Way ANOVA).

