## Supplemental figures 1-10 for "SUMOylation differentially regulates SMCHD1 complex formation and function in a genomic context-specific manner"

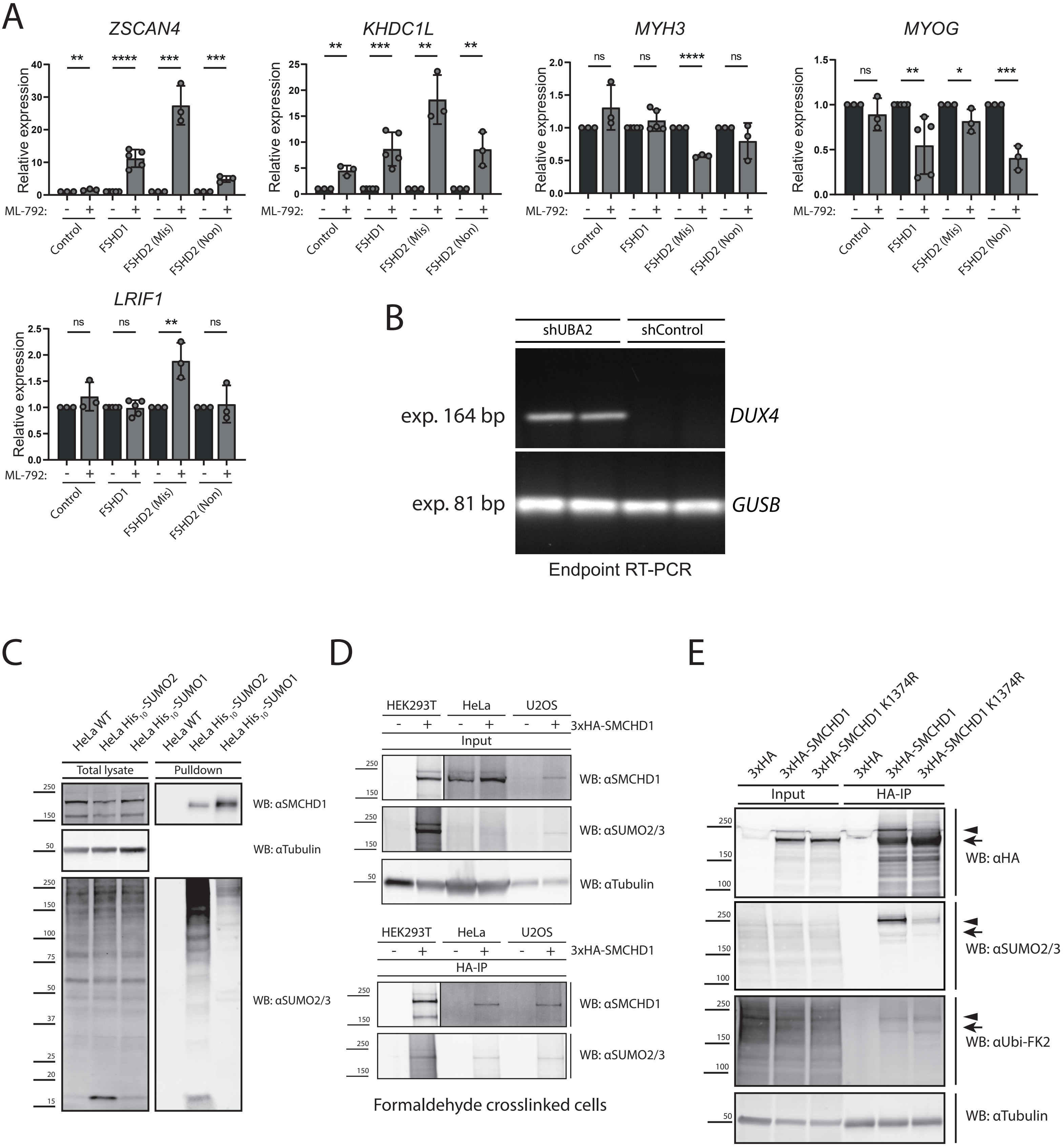
A
ZSCAN4
KHDC1L
MYH3
MYOG
LRIF1
Relative expression
ML-792:
Control
FSHD1
FSHD2 (Mis)
FSHD2 (Non)
ns
*
**
***
****
B
shUBA2
shControl
exp. 164 bp
DUX4
exp. 81 bp
GUSB
Endpoint RT-PCR
C
HeLa WT
HeLa His$_{10}$-SUMO2
HeLa His$_{10}$-SUMO1
Total lysate
Pulldown
WB: αSMCHD1
WB: αTubulin
WB: αSUMO2/3
D
HEK293T
HeLa
U2OS
3xHA-SMCHD1
Input
HA-IP
WB: αSMCHD1
WB: αSUMO2/3
WB: αTubulin
Formaldehyde crosslinked cells
E
3xHA
3xHA-SMCHD1
3xHA-SMCHD1 K1374R
Input
HA-IP
WB: αHA
WB: αSUMO2/3
WB: αUbi-FK2
WB: αTubulin

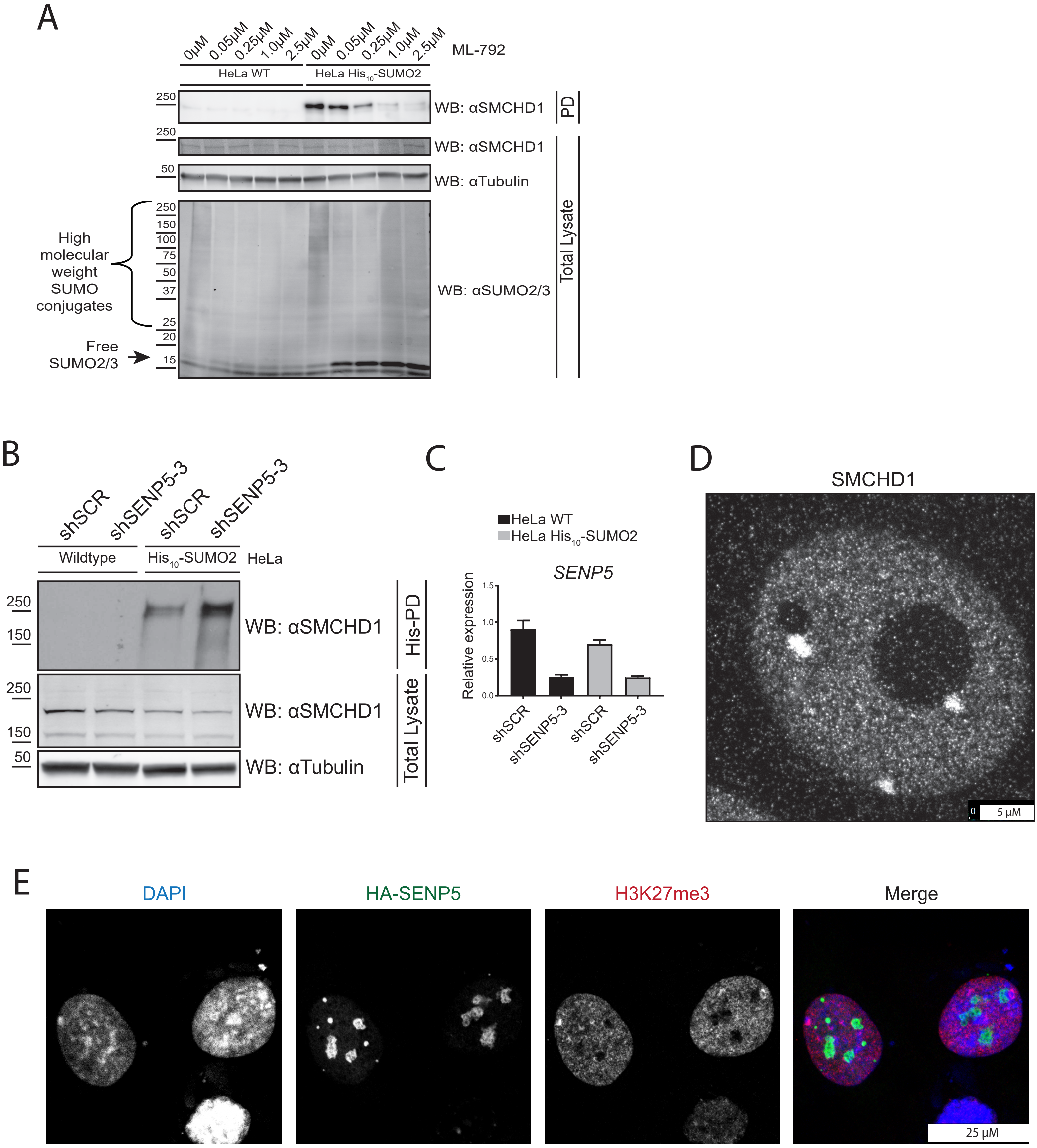
A
0μM 0.05μM 0.25μM 1.0μM 2.5μM 0μM 0.05μM 0.25μM 1.0μM 2.5μM
ML-792
HeLa WT
HeLa His10-SUMO2
250
WB: αSMCHD1
PD
250
WB: αSMCHD1
50
WB: αTubulin
250
150
100
75
50
37
25
20
15
High molecular weight SUMO conjugates
Free SUMO2/3
WB: αSUMO2/3
Total Lysate
B
shSCR
shSENP5-3
shSCR
shSENP5-3
Wildtype
His10-SUMO2
HeLa
250
150
WB: αSMCHD1
His-PD
250
150
WB: αSMCHD1
50
WB: αTubulin
Total Lysate
C
HeLa WT
HeLa His10-SUMO2
SENP5
Relative expression
1.5
1.0
0.5
0.0
shSCR
shSENP5-3
shSCR
shSENP5-3
D
SMCHD1
0
5 μM
E
DAPI
HA-SENP5
H3K27me3
Merge
25 μM

S2

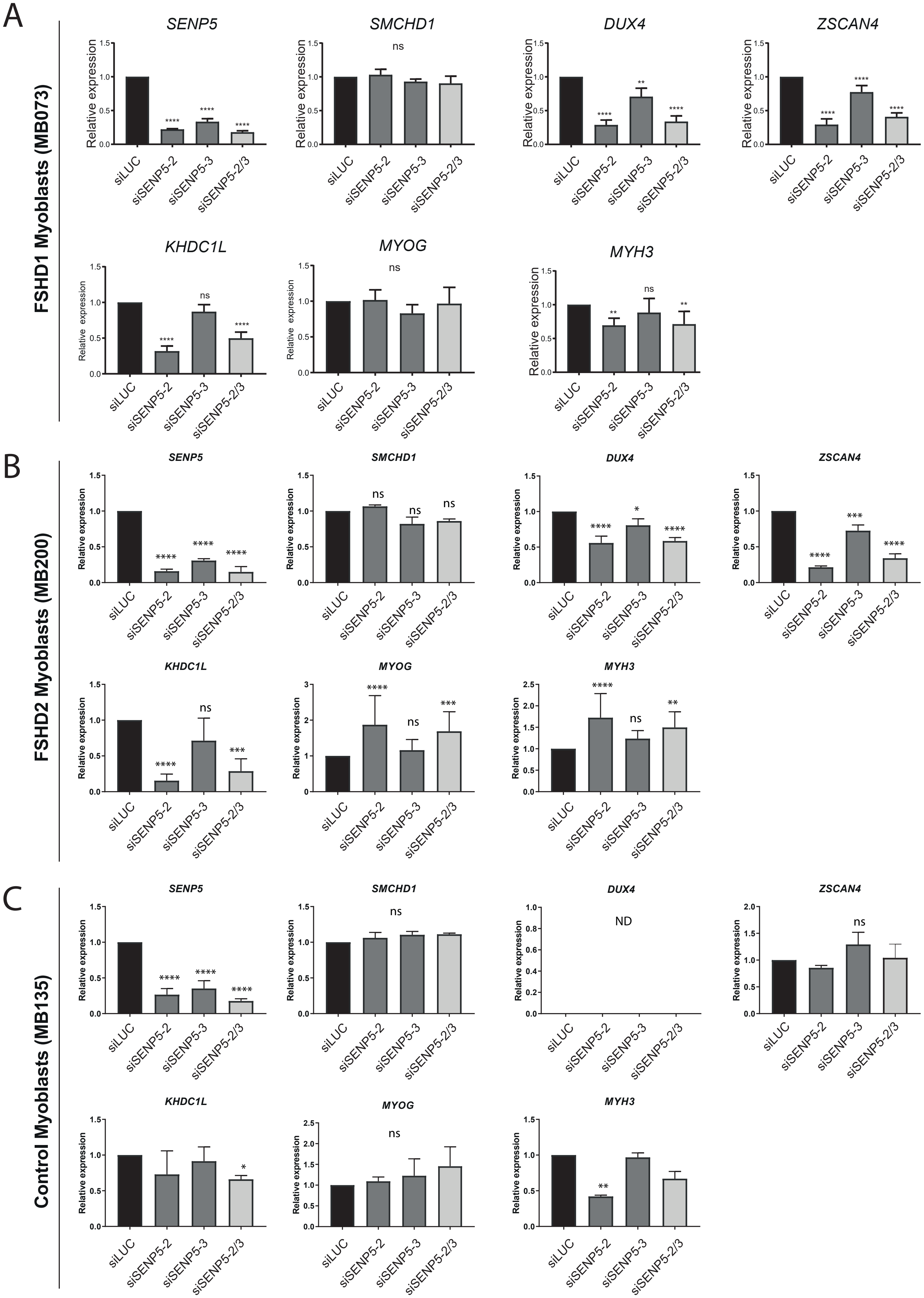
A
FSHD1 Myoblasts (MB073)
SENP5
SMCHD1
DUX4
ZSCAN4
KHDC1L
MYOG
MYH3
Relative expression
siLUC
siSENP5-2
siSENP5-3
siSENP5-2/3
ns
B
FSHD2 Myoblasts (MB200)
C
Control Myoblasts (MB135)
ND

S3

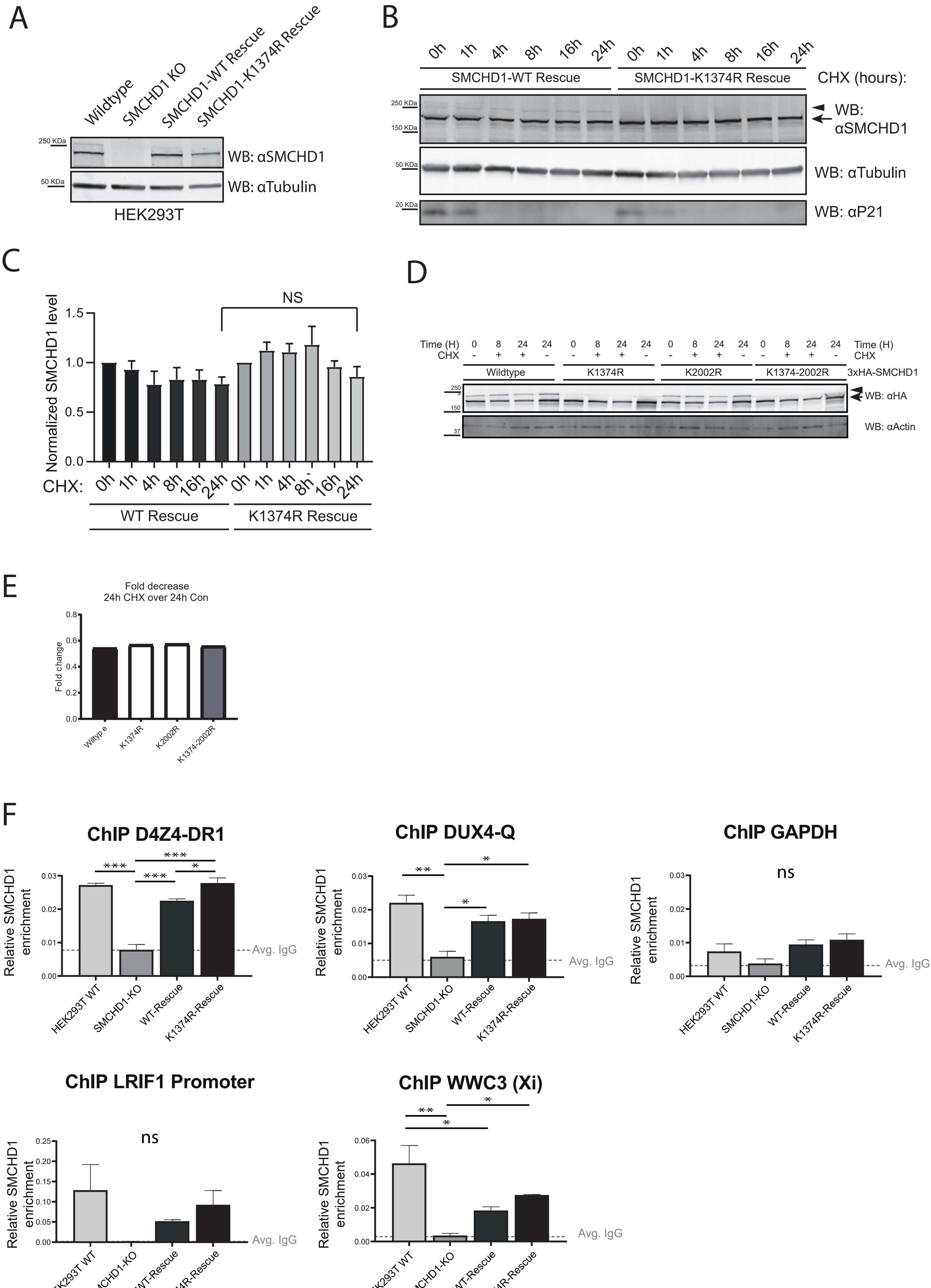
A
Wildtype
SMCHD1 KO
SMCHD1-WT Rescue
SMCHD1-K1374R Rescue
250 KDa
50 KDa
WB: αSMCHD1
WB: αTubulin
HEK293T
B
0h 1h 4h 8h 16h 24h 0h 1h 4h 8h 16h 24h
SMCHD1-WT Rescue
SMCHD1-K1374R Rescue
CHX (hours):
250 KDa
150 KDa
WB: αSMCHD1
50 KDa
WB: αTubulin
20 KDa
WB: αP21
C
NS
Normalized SMCHD1 level
1.5
1.0
0.5
0.0
CHX: 0h 1h 4h 8h 16h 24h 0h 1h 4h 8h 16h 24h
WT Rescue
K1374R Rescue
D
Time (H) 0 8 24 24 0 8 24 24 0 8 24 24 0 8 24 24 Time (H)
CHX - + + - - + + - - + + - - + + - CHX
Wildtype
K1374R
K2002R
K1374-2002R
3xHA-SMCHD1
250
150
WB: αHA
37
WB: αActin
E
Fold decrease
24h CHX over 24h Con
Fold change
0.8
0.6
0.4
0.2
0.0
Wiltype
K1374R
K2002R
K1374-2002R
F
ChIP D4Z4-DR1
ChIP DUX4-Q
ChIP GAPDH
ChIP LRIF1 Promoter
ChIP WWC3 (Xi)
Relative SMCHD1 enrichment
***
**
*
ns
Avg. IgG
HEK293T WT
SMCHD1-KO
WT-Rescue
K1374R-Rescue

S4

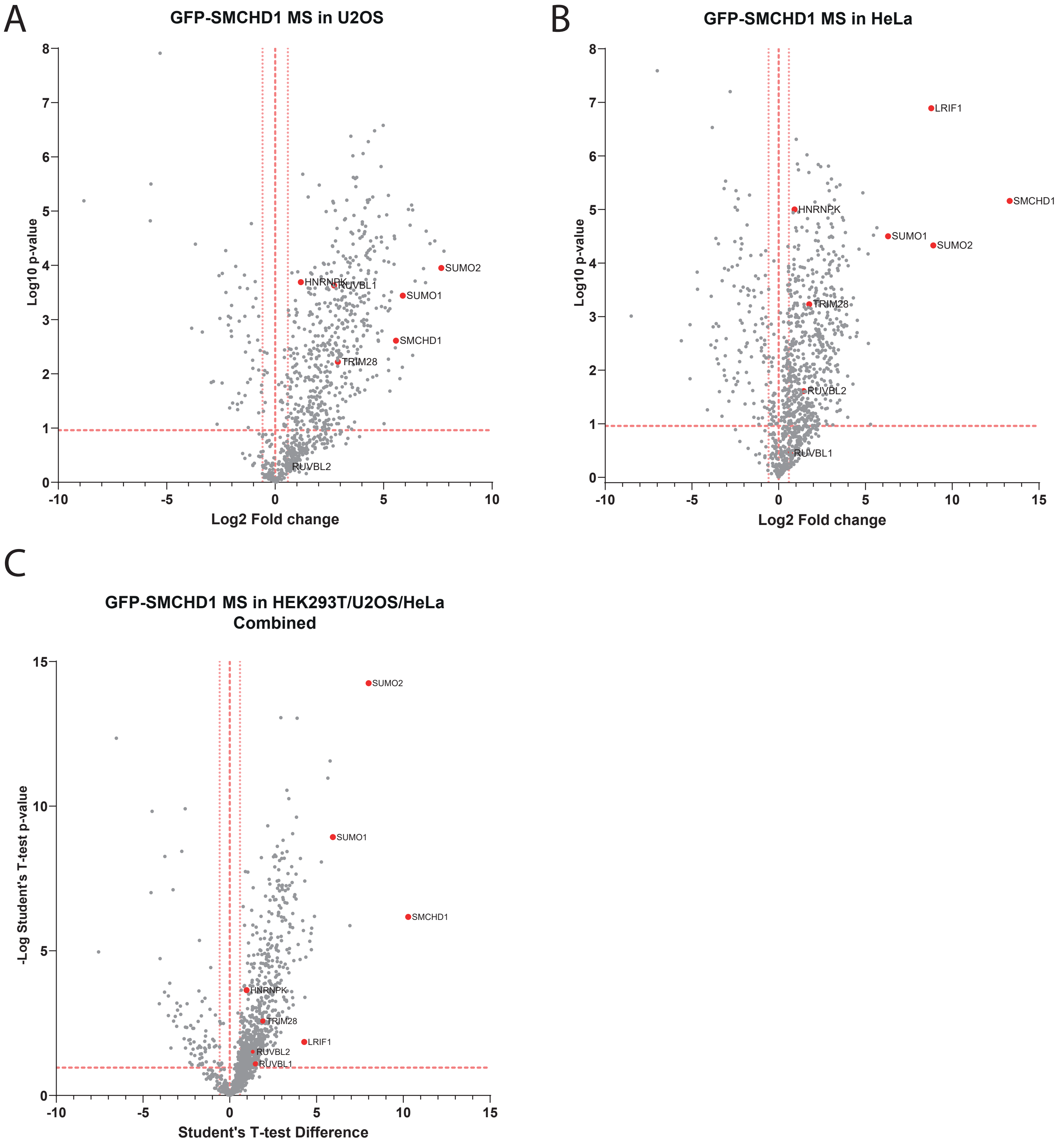
A
GFP-SMCHD1 MS in U2OS
Log10 p-value
Log2 Fold change
SUMO2
HNRNPK
RUVBL1
SUMO1
SMCHD1
TRIM28
RUVBL2
B
GFP-SMCHD1 MS in HeLa
Log10 p-value
Log2 Fold change
LRIF1
SMCHD1
HNRNPK
SUMO1
SUMO2
TRIM28
RUVBL2
RUVBL1
C
GFP-SMCHD1 MS in HEK293T/U2OS/HeLa
Combined
-Log Student's T-test p-value
Student's T-test Difference
SUMO2
SUMO1
SMCHD1
HNRNPK
TRIM28
LRIF1
RUVBL2
RUVBL1

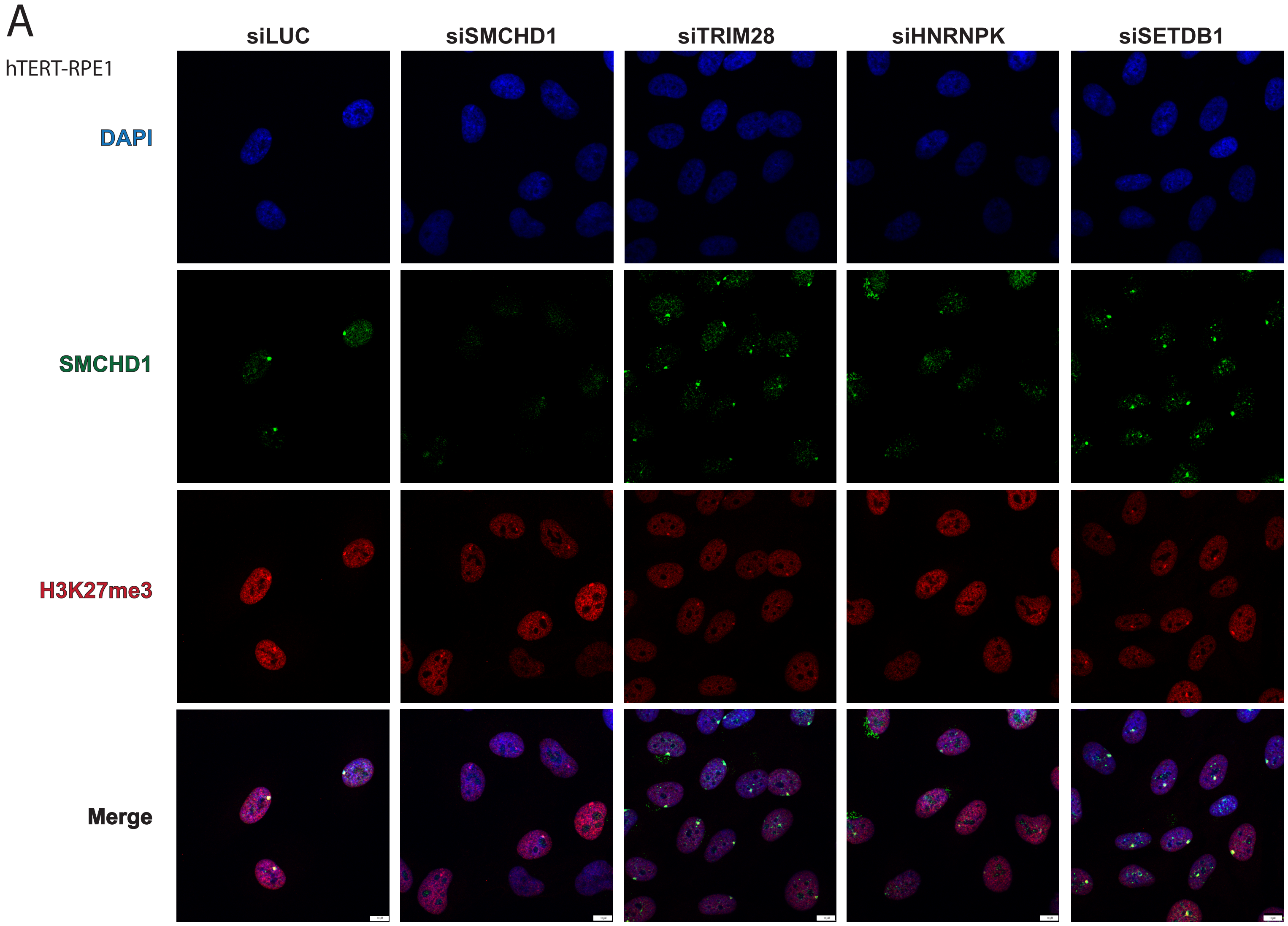
A
hTERT-RPE1
siLUC
siSMCHD1
siTRIM28
siHNRNPK
siSETDB1
DAPI
SMCHD1
H3K27me3
Merge

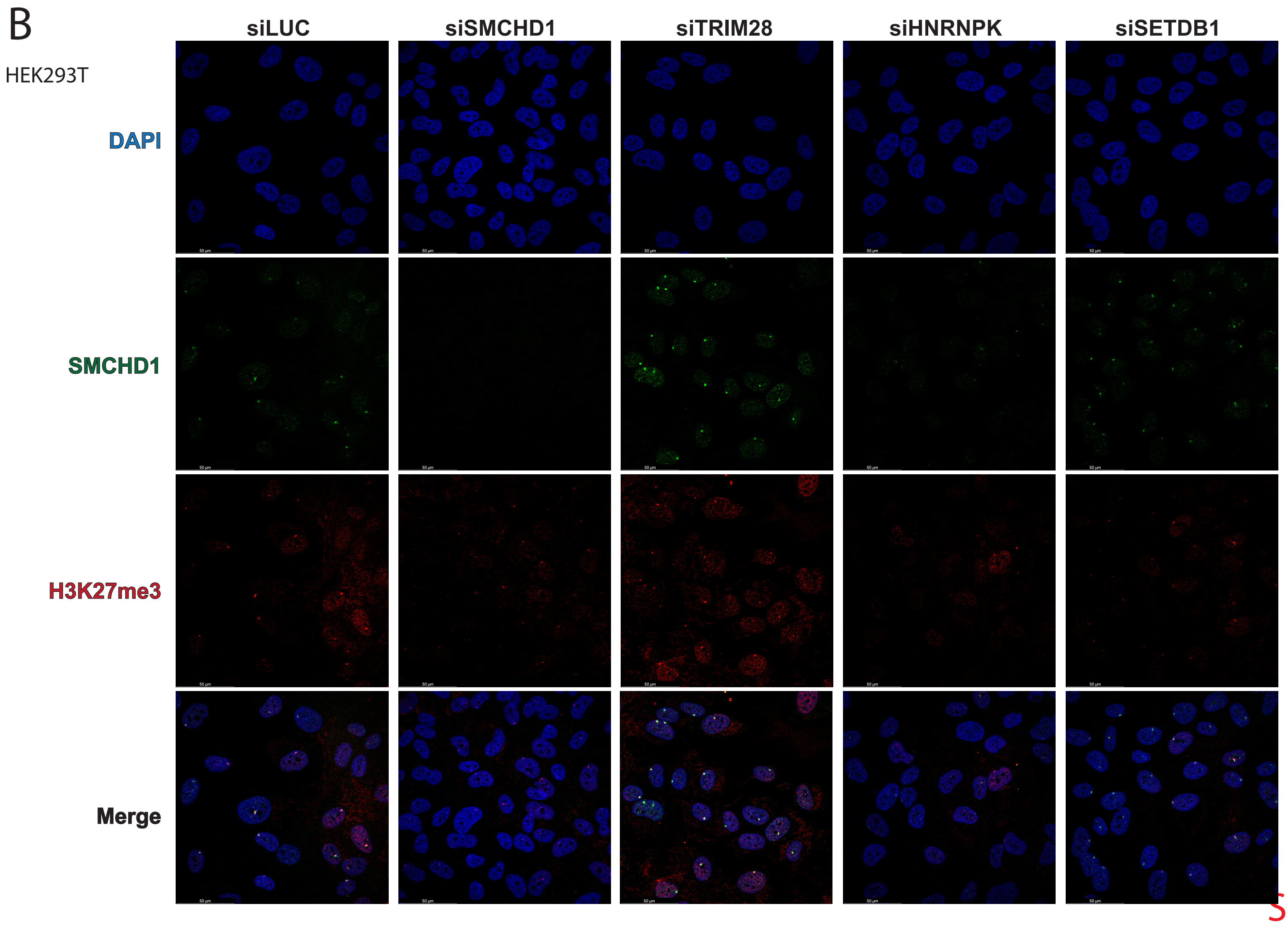
B
HEK293T
siLUC
siSMCHD1
siTRIM28
siHNRNPK
siSETDB1
DAPI
SMCHD1
H3K27me3
Merge

A

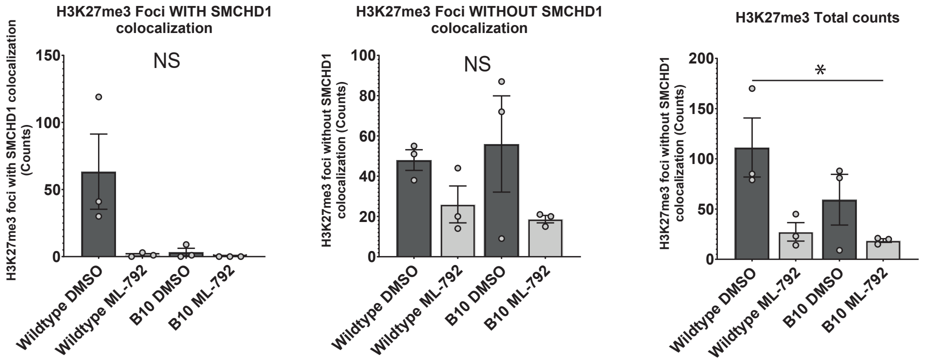


B

| H3K27me3 foci **with** SMCHD1 colocalization | H3K27me3 foci **without** SMCHD1 colocalization | **Total** H3K27me3 foci | Total nuclei | Percent of H3K27me3 foci with SMCHD1 colocalization | **Sample:** |
|---|---|---|---|---|---|
| 190 | 144 | 334 | 285 | 57% | **WT DMSO** |
| 4 | 78 | 82 | 244 | 5% | **WT ML-792** |
| 10 | 168 | 178 | 249 | 6% | **KO DMSO** |
| 0 | 56 | 56 | 166 | 0% | **KO ML-792** |

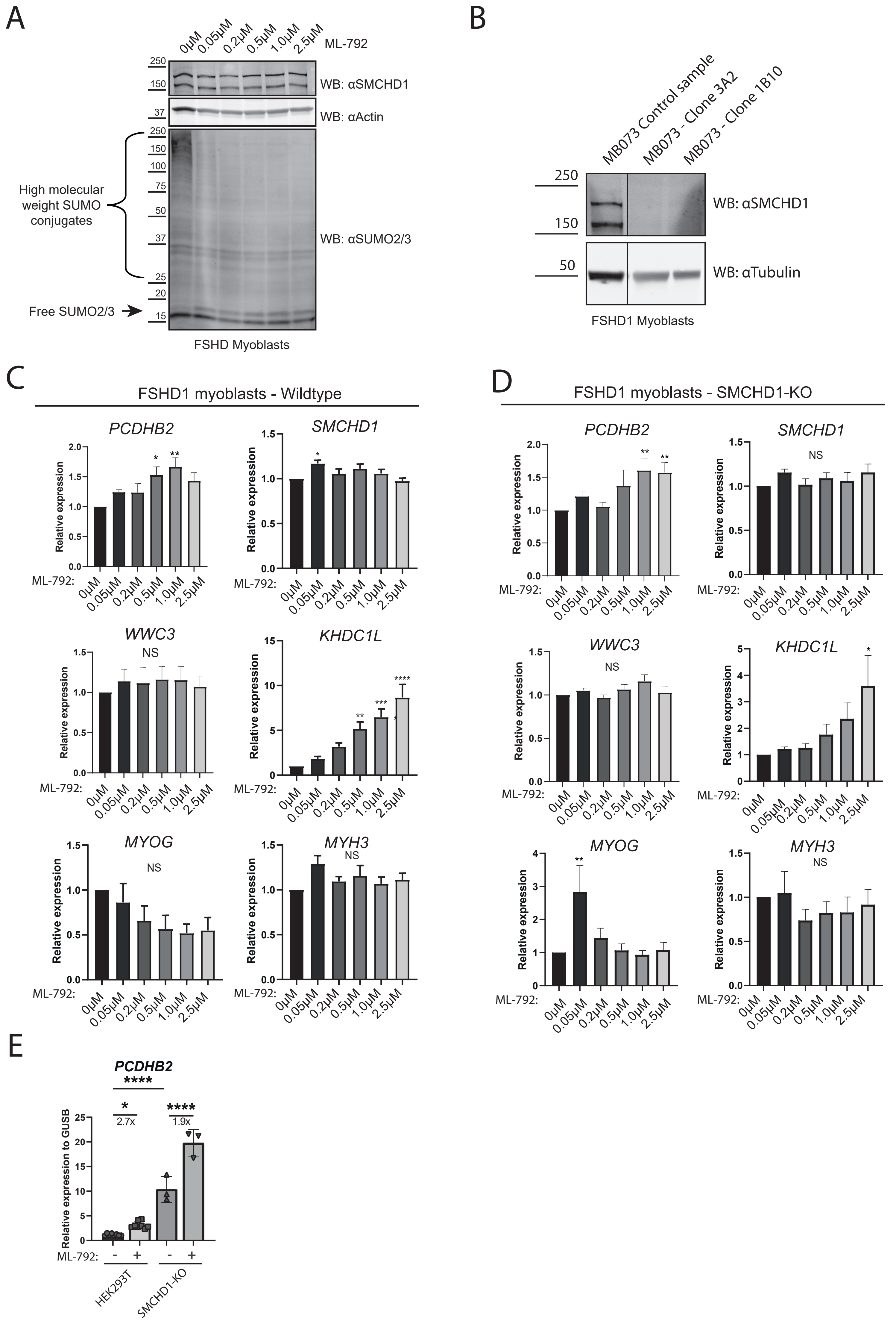
A
0µM 0.05µM 0.2µM 0.5µM 1.0µM 2.5µM ML-792
WB: αSMCHD1
WB: αActin
High molecular weight SUMO conjugates
Free SUMO2/3
WB: αSUMO2/3
FSHD Myoblasts
B
MB073 Control sample
MB073 - Clone 3A2
MB073 - Clone 1B10
WB: αSMCHD1
WB: αTubulin
FSHD1 Myoblasts
C
FSHD1 myoblasts - Wildtype
PCDHB2
SMCHD1
WWC3
KHDC1L
MYOG
MYH3
NS
Relative expression
ML-792:
D
FSHD1 myoblasts - SMCHD1-KO
PCDHB2
SMCHD1
WWC3
KHDC1L
MYOG
MYH3
NS
Relative expression
ML-792:
E
PCDHB2
2.7x
1.9x
Relative expression to GUSB
ML-792:
HEK293T
SMCHD1-KO

S8

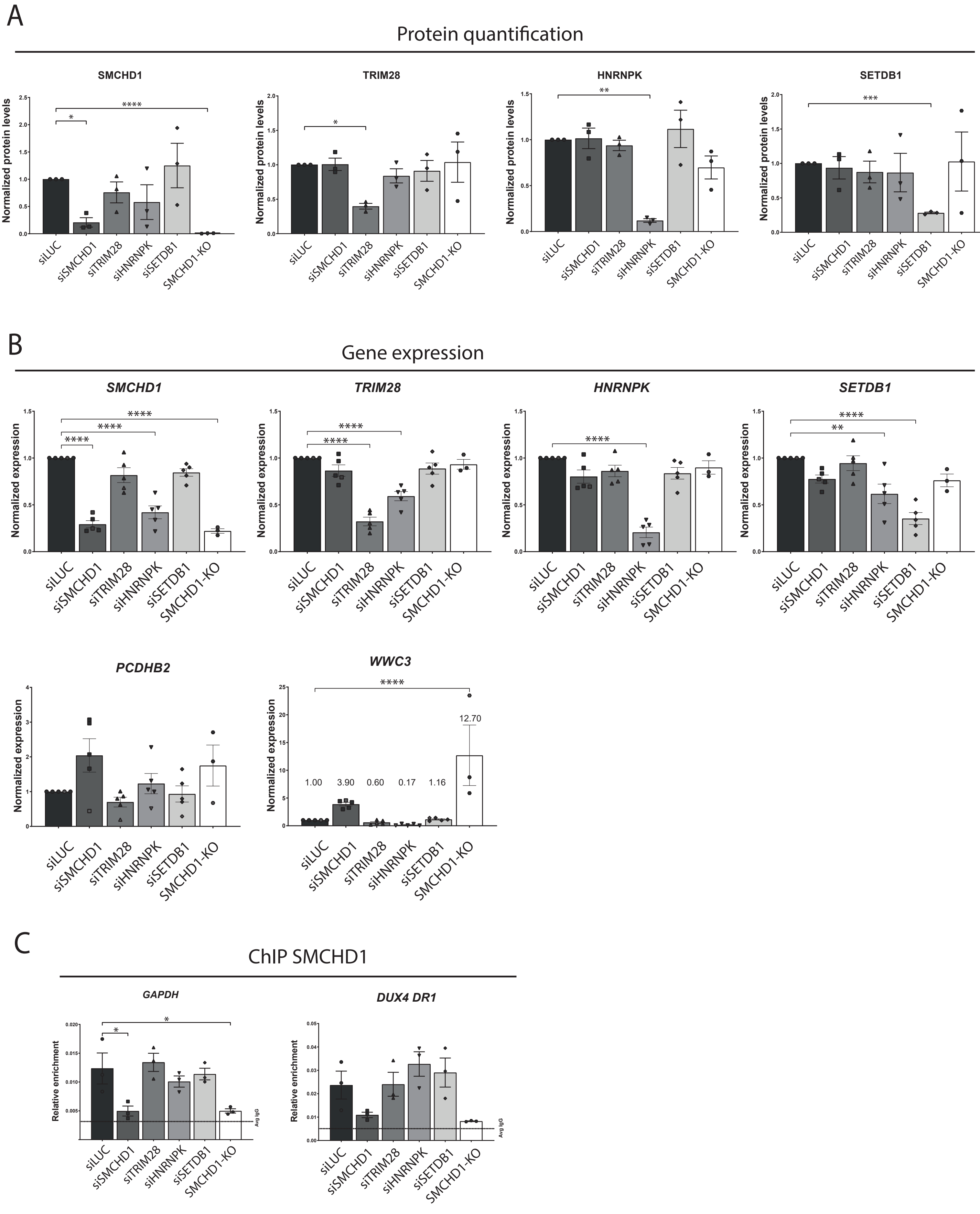
A
Protein quantification
SMCHD1
TRIM28
HNRNPK
SETDB1
Normalized protein levels
siLUC
siSMCHD1
siTRIM28
siHNRNPK
siSETDB1
SMCHD1-KO
B
Gene expression
SMCHD1
TRIM28
HNRNPK
SETDB1
PCDHB2
WWC3
Normalized expression
1.00
3.90
0.60
0.17
1.16
12.70
C
ChIP SMCHD1
GAPDH
DUX4 DR1
Relative enrichment
Avg IgG

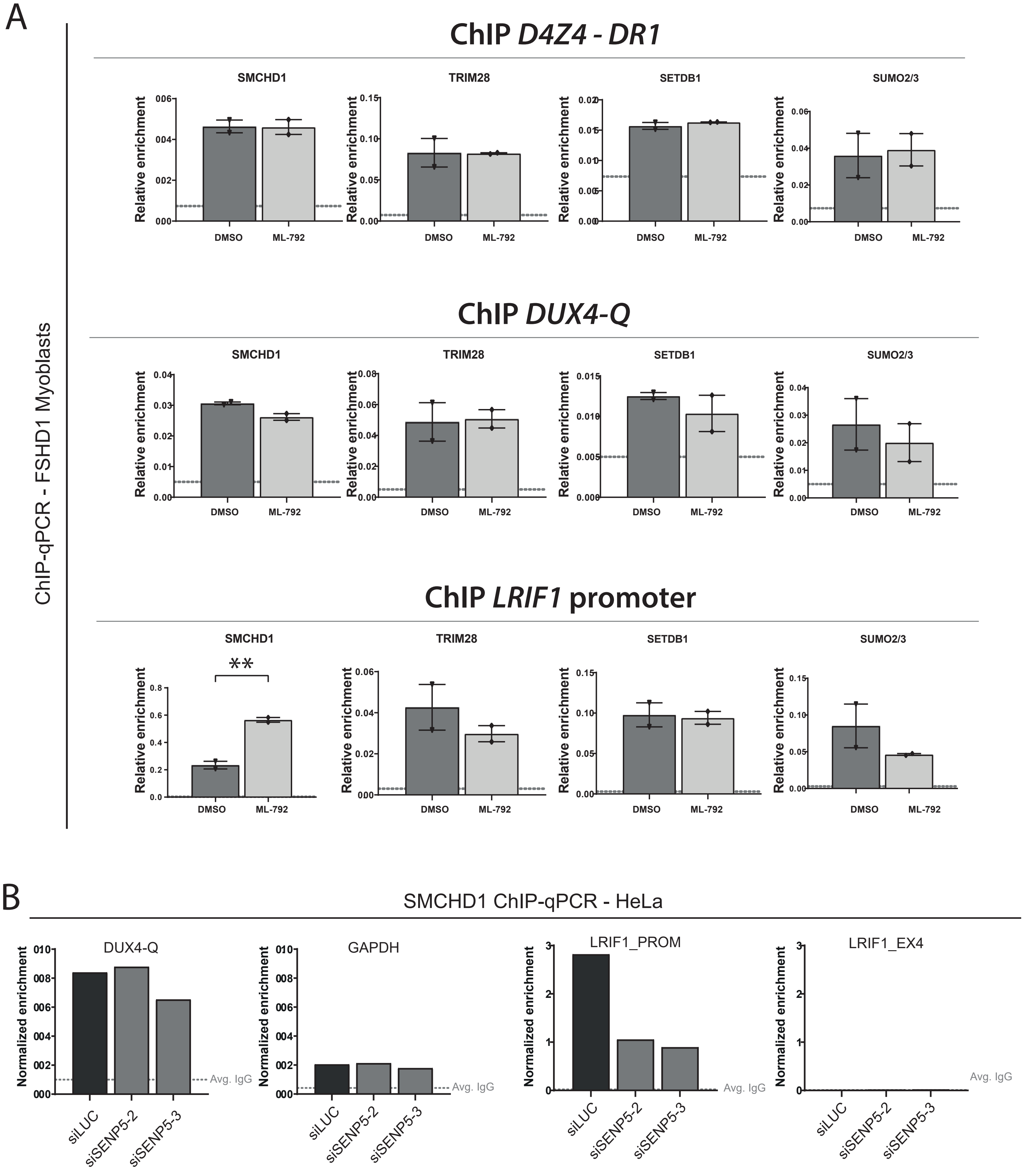
A
ChIP-qPCR - FSHD1 Myoblasts
ChIP D4Z4 - DR1
ChIP DUX4-Q
ChIP LRIF1 promoter
SMCHD1
TRIM28
SETDB1
SUMO2/3
Relative enrichment
DMSO
ML-792
**
B
SMCHD1 ChIP-qPCR - HeLa
DUX4-Q
GAPDH
LRIF1_PROM
LRIF1_EX4
Normalized enrichment
Avg. IgG
siLUC
siSENP5-2
siSENP5-3
